# Direct effector recognition by a tandem kinase triggers non-canonical immunity in wheat

**DOI:** 10.64898/2026.02.17.706082

**Authors:** Caitlin Lemmer, Emmanuel N. Annan, Benjamin Schwessinger, Eric Pereira, Bayantes Dagvadorj, Di Xiao, Brande B. H. Wulff, Simon G. Krattinger, Ann Kwan, Jameel M. Abduljalil, Smriti Singh, Thuan Nha Ho, Brent N. Kaiser, Evans Lagudah, Robert F. Park, Peng Zhang, Li Huang, Yi Ding

## Abstract

Tandem kinase proteins (TKPs) are an important class of disease resistance genes in cereal crops alongside nucleotide-binding (NB-ARC) leucine-rich repeat (LRR) (NLR) receptors. Here, we cloned the wheat (*Triticum aestivum*) resistance gene *Lr41*, which encodes a TKP with a C-terminal fusion, and its paired effector gene *AvrLr41* from the rust pathogen *Puccinia triticina*. AvrLr41 is unusually large and highly divergent from most fungal effectors and is directly recognized by Lr41 through its pseudokinase domain. Immune activation mediated by Lr41 require auxiliary helper proteins, including either a minimal coiled-coil (CC) helper protein or its full-length NLR ortholog. These findings reveal a non-canonical immune system in which a kinase-type receptor directly recognizes an effector and recruits either CC or NLR helpers, demonstrating that full-length NLRs are not strictly required for NLR-linked immune execution and expanding the functional repertoire of disease resistance proteins in crops.

## Main

The deployment of host resistance (*R*) genes is a sustainable strategy for managing crop diseases. Many *R* genes cloned from plants to date encode intracellular nucleotide-binding leucine-rich repeat (NLR) receptors that recognize pathogen-secreted effectors [avirulence proteins (Avr)] and activate immune signaling^1^. Avr recognition by NLRs may be direct or indirect^2,3^, converging on conserved outputs such as resistosome or hetero-oligomer assembly, calcium-permeable channel activity, and hypersensitive cell death^4–10^. Indirect interactions often involve kinases or pseudokinases partners, such as the tomato protein kinase Pto guarded by Prf^11^, kinase PBS1 monitored by RPS5^12^, and the recognition of pseudokinase ZED1 or RKS1 by ZAR1^13–15^.

In Triticeae, an alternate class of *R* gene that accounts for ∼20% cloned wheat and barley *R* genes^16^ encodes tandem kinase proteins (TKPs). Since the first identified barley stem rust resistance protein (Rpg1)^17^, TKPs have been shown to confer resistance to diverse wheat fungal pathogens, including the stripe rust pathogen *Puccinia striiformis* f. sp. *tritici* (*Pst*; Yr15/WTK1)^18^, the stem rust pathogen *P. graminis* f. sp. *tritici* (*Pgt*; Sr60/WTK2, Sr62/WTK5)^19,20^, the leaf rust pathogen *P. triticina* (Lr9/WTK6-vWA, Lr39/WTK8-msp-WD40)^21,22^, the powdery mildew pathogen *Blumeria graminis* f. sp. *tritici* (Pm24/WTK3, Pm36/WTK7-TM, Pm57/WTK6b-vWA)^23–25^ and the wheat blast pathogen *Magnaporthe oryzae* (Rwt4)^26^. Although TKP models were hypothesized to be analogous to Pto/RKS1/ZED1^27^, recent studies revealed direct sensing by Sr62 and WTK3/Rwt4 of their respective effectors AvrSr62 and PWT4 via their N-terminal kinase domains, with the C-terminal domains relaying signal to the same helper NLR, wheat tandem NBD 1 (WTN1)^28,29^. The proposed modes of action of Sr62 and WTK3 are similar, with WTK3-WTN1 facilitating formation of calcium-permeable channels. Whether helper NLRs are a general requirement for TKP-mediated immunity remains unknown. Unlike Sr62 and WTK3, many identified TKPs harbor diverse C-terminal domain fusions with functions in immunity that remain unclear.

Research on cereal rust resistance has been hampered because the obligate biotrophic lifestyles and highly heterozygous genomes of rust fungi make it difficult to identify their *Avr* genes. To date, fewer than 10 *Avr* genes have been conclusively identified, predominantly from *Pgt*^28,30–35^. These Avr proteins are generally compact, lack sequence homology, and bind directly to their corresponding NLR receptors. Among them, AvrSr35 is unusually large in size^32^, and AvrSr62 directly binds to the TKP member Sr62^28^. Although candidate Avrs have been described in *P. triticina*^36^, no Avr-R interaction model is firmly established for this species complex, despite leaf rust being a major threat to global wheat production^37,38,39^ and the most damaging of all wheat pathogens and pests^1^.

Here, we cloned the wheat leaf rust resistance gene *Lr41*, a TKP derived from the wheat wild relative *Aegilops tauschii*, and identified its corresponding effector *AvrLr41* in *P. triticina.* In addition, we identified two helper proteins, including an unconventional protein truncated from an NLR at the *Lr41* locus lacking NB-ARC and LRR domains and its full-length ortholog from *Ae. tauschii*, each of which can cooperate with Lr41 and AvrLr41 to trigger the hypersensitive response (HR). This adds a model distinct from the prevailing paradigm in which full-length NLRs are required for immune signaling, and suggests an alternative recognition mechanism with implications for broadening our understanding of NLR functional architectures.

## Results

### *Lr41* encodes a tandem kinase with C-terminal fusions

*Lr41* confers resistance to wheat leaf rust pathogen at all growth stages. It was initially mapped on chromosome arm 1DS and later relocated to 2DS in wheat. The source of *Lr41* was reported to be the *Ae. tauschii* accession TA1675 on 2DS based on analysis with telosomic and single sequence repeat (SSR) markers^40,41^. *Xbarc124* is the closest marker, located about 1 cM from *Lr41*^42^. We inspected the TA1675 contigs^22^ for the presence of three markers linked to *Lr41*^42^. We then aligned the TA1675 and Chinese Spring (CSv2.0) sequences from their respective 2DS arms (Fig. 1a-b), which allowed us to narrow down the distal chromosome region containing *Lr41* to a 5.7-Mb interval in the TA1675 genome (Fig. 1c). Using mRNA-seq data, we found no candidate full-length NLR gene with polymorphisms among *T. aestivum* and *Ae. tauschii* lines carrying or lacking *Lr41* (table S1-2; Fig. S3). We then analysed transcriptome and genome sequence data from +*Lr41* and -*Lr41* lines, along with a panel of *Ae. tauschii* lines known to lack *Lr41*, and carried out syntenic analysis between the distal ends of the 2DS arms from TA1675 and CS. By refining new gene content potentially introgressed into the Thatcher (Tc) near-isogenic line (NIL) Tc+*Lr41* over the 5.7-Mb interval, all mRNA-seq reads were remapped and quantified for transcript abundance. We identified 74 intervals with transcripts exclusively in the Tc+*Lr41* line and TA1675 and absent in Tc (Fig. 1d-e). Among these transcripts, two intervals contained single, intact gene models producing full-length transcripts with moderate expression levels in Tc+*Lr41* (Fig. 1d-e; Fig. S2a). No synteny was observed between TA1675 and CS for these two intervals (Fig. 1d). The full-length transcripts carried nonsense mutations in all *Ae. tauschii* lines lacking *Lr41* (Fig. S2a).

**Fig. 1.**
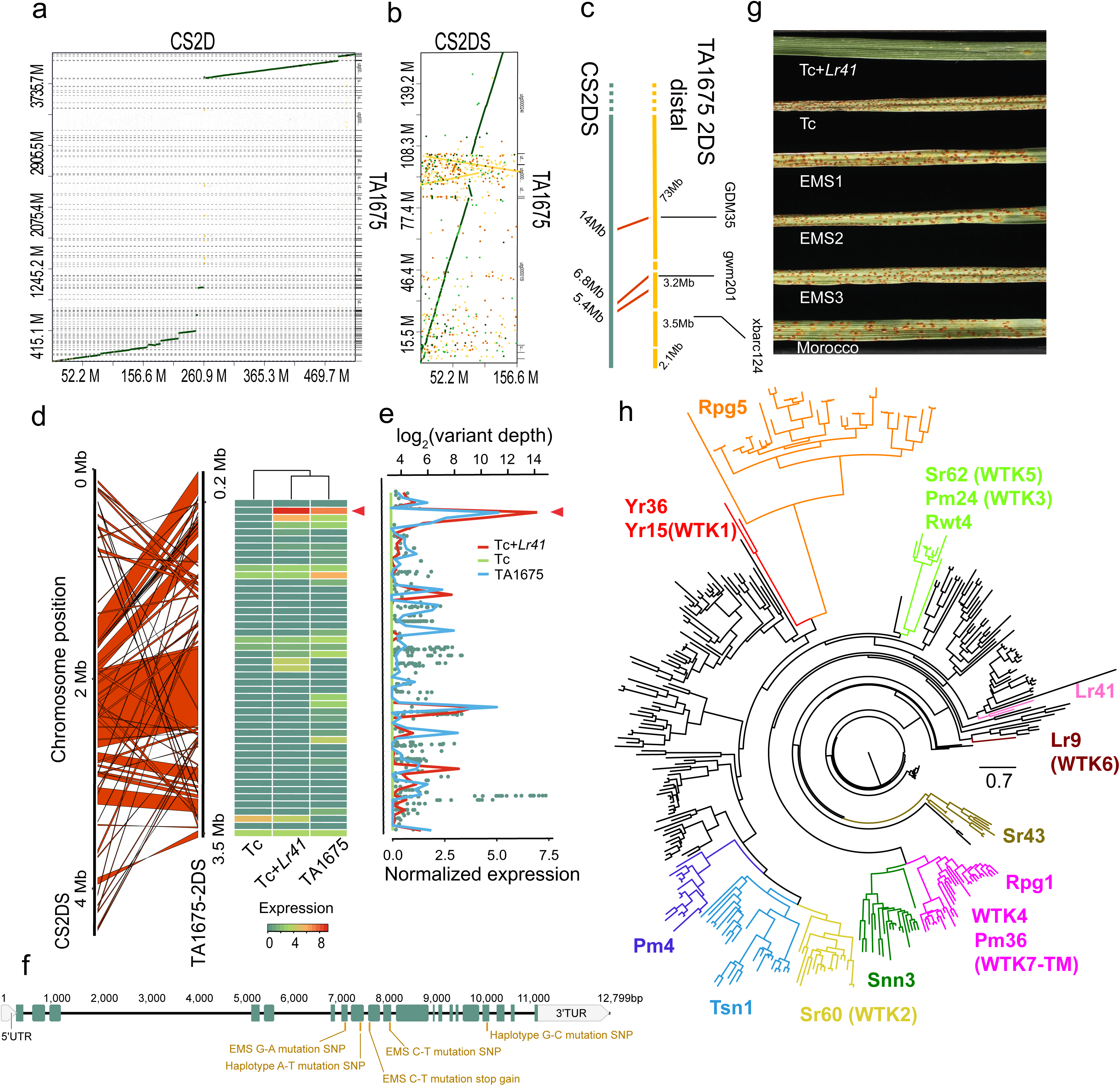
Genome-assisted identification of *Lr41*. (**a**) Homology-based scaffolding of contigs of the *Aegilops tauschii* line TA1675 aligned to chromosome 2D of the Chinese Spring (CS) reference genome. (**b**) Zoomed-in view of an alignment between the CS 2D short arm (2DS) genomic region and TA1675 contigs. Diagonal lines indicate conserved syntenic segments. Dark green, light green to orange and yellow represent high to low homologies. (**c**) Physical positions of markers define TA1675 contigs comprising the distal 2DS interval containing *Lr41*. (**d**) Synteny of the ∼3.5 Mb distal 2DS region of TA1675 with CS 2DS. Red bands indicate syntenic regions. The accompanying heatmap shows normalized FPKM expression values for transcripts under multiple sample conditions. Transcripts were labelled as coordinate regions. (**e**) Expression pattern and sequence variation across the TA1675 contigs corresponding to the 2DS-assigned region. The line plot depicts normalized transcript abundance, and overlaid dots represent homozygous variant sites with variant depth normalized. Arrows denote Lr41 coding regions. (**f**) Gene model of *Lr41*, comprising 21 exons spanning ∼12.8 kb, including 5’ and 3’ untranslated regions (UTRs). EMS-induced mutations detected in mutants and single nucleotide polymorphisms (SNPs) detected in additional *Ae. tauschii* lines are indicated. (**g**) Representative infection phenotypes of Tc+*Lr41*, Tc, three EMS-derived mutants, and Morocco control following inoculation with *Puccinia triticina* (pathotype 64-11). (**h**) Maximum-likelihood phylogeny of 369 cereal kinase-type resistance proteins, including tandem-kinase proteins (TKPs) and related kinase families. Lr41 clusters within a distinct branch separate from other TKPs such as Sr62 (WTK5), Rwt4/Pm24 (WTK3), Yr15 (WTK1), Lr9 (WTK6), Pm36 (WTK7-TM), and Sr60 (WTK2). The scale bar indicates substitutions per site.

The two candidate intervals together correspond to a single coding region that encodes a full-length protein comprising two tandem kinase domains, followed by a Major Sperm Protein (MSP) domain and WD40 repeats (Fig. S2b). In a complementary approach, we screened ∼4,000 ML plants from EMS-treated Tc+*Lr41* lines using an avirulent *P. triticina* pathotype 64-11 (s473) and identified three mutants (Fig. S3; table S1-2), where we confirmed non-synonymous transition mutations in this coding region: G->A (+ 1100), C->T (+1459, stop-gain), and C->T (+1793), as validated by amplicon sequencing (Fig. 1f-g and S4). Additional functional validation through virus-induced gene silencing (VIGS) of the candidate gene in Tc+*Lr41* plants using an avirulent North American pathotype PBJJG resulted in diminished resistance (Fig. S1a-b). On this basis, we considered the full-length candidate gene to be *Lr41*.

*Lr39* and *Lr41* have been long hypothesized to be closely linked or allelic^40,41^. We found the *Lr41* sequence to be identical to the recently cloned *Lr39*^22^, consistent with the shared virulence profiles of their respective NILs (Fig. S1c), thereby confirming that *Lr41* and *Lr39* represent the same gene. Lr41/Lr39 is distinct from other known kinase-type R proteins, as supported by a phylogenetic tree constructed from 369 cereal kinases from various R families (Fig. 1h). Two haplotype variations were previously identified in *Lr39*, including a substitution of a positively charged amino acid with a hydrophobic residue (R457I) in the kinase 2 (K2) domain^22^. In contrast, the current EMS-induced SNPs both occur in the K2 coding region, introducing a premature stop codon and non-synonymous mutations that lead to amino acid (aa) changes predicted to disrupt hydrogen-bond geometry (S367N) and cause steric interference (S598F) (Fig. S2b). These alterations likely compromise the structural integrity and functional capacity of the K2 domain, which is essential for Lr41-mediated resistance.

### A highly variant region in *P. triticina* is associated with pathogenicity to *Lr41*

*P. triticina* is dikaryotic, containing two genetically distinct and often highly heterozygous haploid nuclei (haplotype 1 and haplotype 2). In this scenario, spontaneous gain of virulence to a single resistance gene may arise from mutations in the corresponding avirulence locus in one haplotype. To systematically associate genomic variation with *Lr41* virulence, we performed genome-wide association study (GWAS) using genome and RNA-seq data from 52 *P. triticina* isolates and their corresponding phenotypes (NCBI accession PRJNA1394492; table S3), using the haplotype-resolved reference genome of isolate s473^59^. We included two independent spontaneous *P. triticina* mutants (s543 and s692) that acquired virulence specifically to *Lr41* under greenhouse conditions in the GWAS, together with their respective *Lr41*-avirulent parental isolates (s473 and s365; table S3). Strikingly, highly significant associations by GWAS for *Lr41* virulence were detected only with the haplotype 2 reference genome, whereas no comparable signal was observed using the haplotype 1 reference (Fig. 2d; Fig. S5a-c; table S4). Consistent with this haplotype-specific association, alignment of sequencing reads to the s473 haplotype 2 genome revealed a ∼1.3-Mb region uniquely enriched for homozygous SNPs in the two *Lr41*-virulent mutants, but not in *Lr41*-avirulent isolates (s365, s423, and s625), while overall variant site depth remained comparable across isolates (Fig. 2a-c). Notably, this region of lower heterozygosity precisely coincided with the GWAS peak observed in haplotype 2. A total of 284 genes were annotated within this interval (table S5), among which the two most significant nonsynonymous SNPs from GWAS mapped to a single candidate effector gene (Fig. 5d; table S4-5), strongly implicating this gene as *AvrLr41*.

**Fig. 2.**
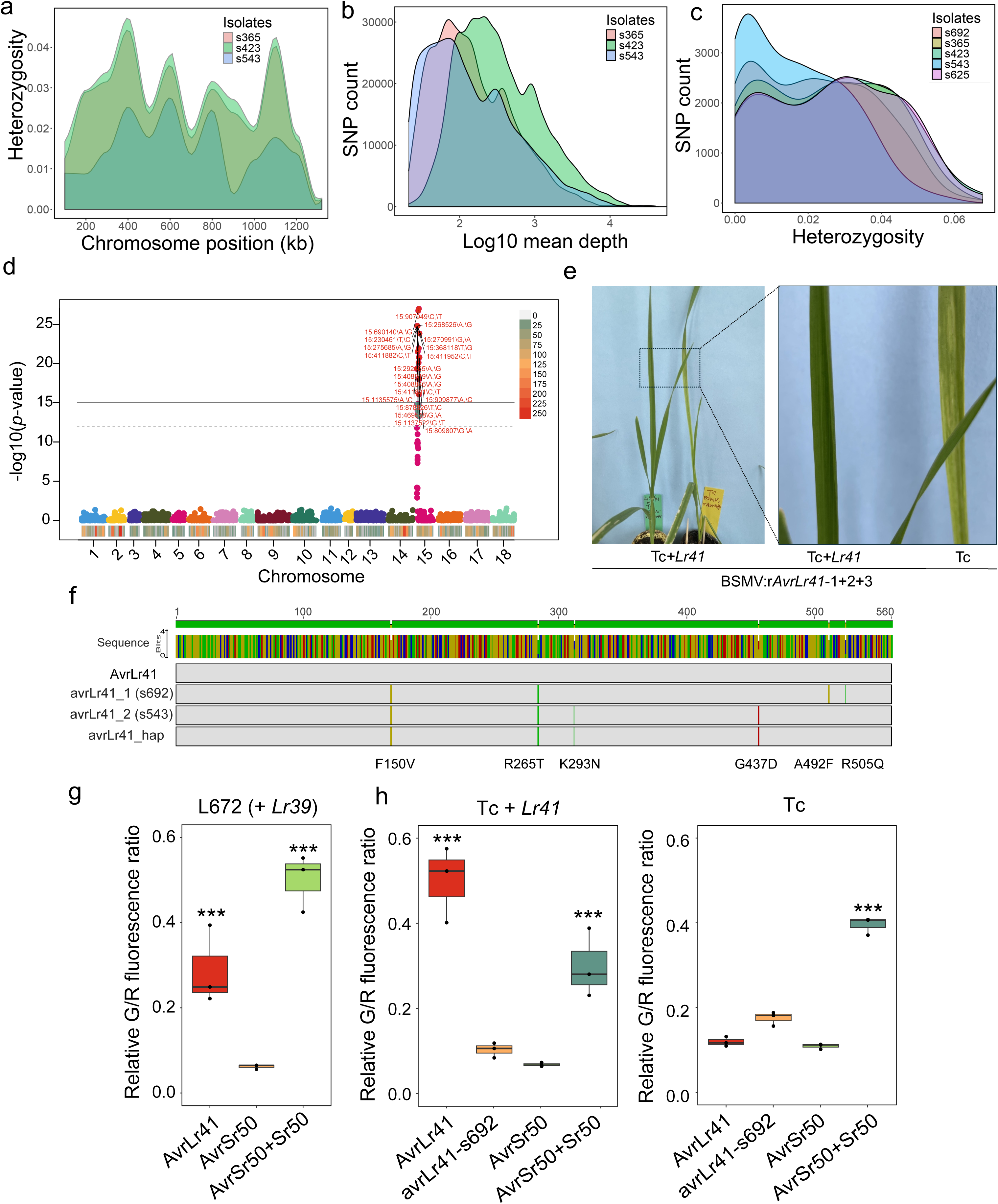
Discovery and functional characterization of AvrLr41. **(a)** Overview of the heterozygosity landscape across a ∼1.3-Mb chromosomal region enriched with homozygous variants in a spontaneous mutant s543, which gained virulence on *Lr41*, compared with avirulent isolates, including its parent s423. (**b**) Distribution of mean SNP depth across isolates over the same genomic interval, illustrating distribution of SNP numbers over calling coverage. (**c**) Heterozygosity profiles for five *Puccinia triticina* isolates, highlighting the lower heterozygosity signal in the *Lr41*-virulent mutants s543 and s692. Heterozygosity ratios were assessed in 100-kb windows. (**d**) Haplotype-based genome-wide association study (GWAS). Two horizontal lines mark the genome-wide significance thresholds at *p* = 1×10L¹² and *p* = 1×10^-15^. The color key represents local SNP density (1-Mb bins) across the genome. (**e**) Assessment of a candidate gene for *AvrLr41* by barley stripe mosaic virus (BSMV)-mediated overexpression, based on the identification of two top SNPs (907,949 bp and 909,877 bp) in the *P. triticina* genome. Three overlapping fragments (*AvrLr41*-1+2+3) were co-expressed. Representative wheat plants are shown at 9 dpi from 20 plants. Three independent experiments were performed with similar results. (**f**) Sequence alignment of AvrLr41 with alleles avrLr41_1 and avrLr41_2 from the virulent isolates s542 and s692 and avrLr41_hap from the avirulent reference isolate s473. Nonsynonymous substitutions distinguishing AvrLr41 and avrLr41 variants include F150V, R265T, K293N, G437D, A492F, and R505Q (positions based on removal of the 19-aa signal peptide). (**g**) Wheat protoplast assay in synthetic hexaploid L672 carrying *Lr39* showing strong defense activation by AvrLr41 but not by the AvrSr50 control. (**h**) Wheat protoplast assays in the Tc+*Lr41* and Tc backgrounds. Co-expression of *AvrSr50* and *Sr50* was used as a positive control. The experiments were repeated three times with similar results. Asterisks indicate significant differences relative to the AvrSr50 negative control, based on Student’s *t*-test (***, *p* < 0.01).

The top candidate gene for *AvrLr41* contains 24 exons (Fig. S9a) and encodes a 560-aa protein (Fig. 2e) with a predicted 20-aa N-terminal signal peptide. It shows no sequence homology to genes in any other species and exhibits low-confidence structural predictions by AlphaFold3 (AF3) (Fig. 3e; table S6). Analysis of sequence variation across *AvrLr41* revealed distinct haplotypes associated with avirulent and virulent phenotypes (Fig. S9a). The avirulent isolate s473 carries a heterozygous F169V substitution (T505G), whereas virulent mutants encode alternative haplotypes with aa changes, including R284T (G851C) in both mutants, K312N (G936T) in s543, and either A511P (G1531C) or R524C (G1571A) in s692 (Fig. 2e).

**Fig. 3.**
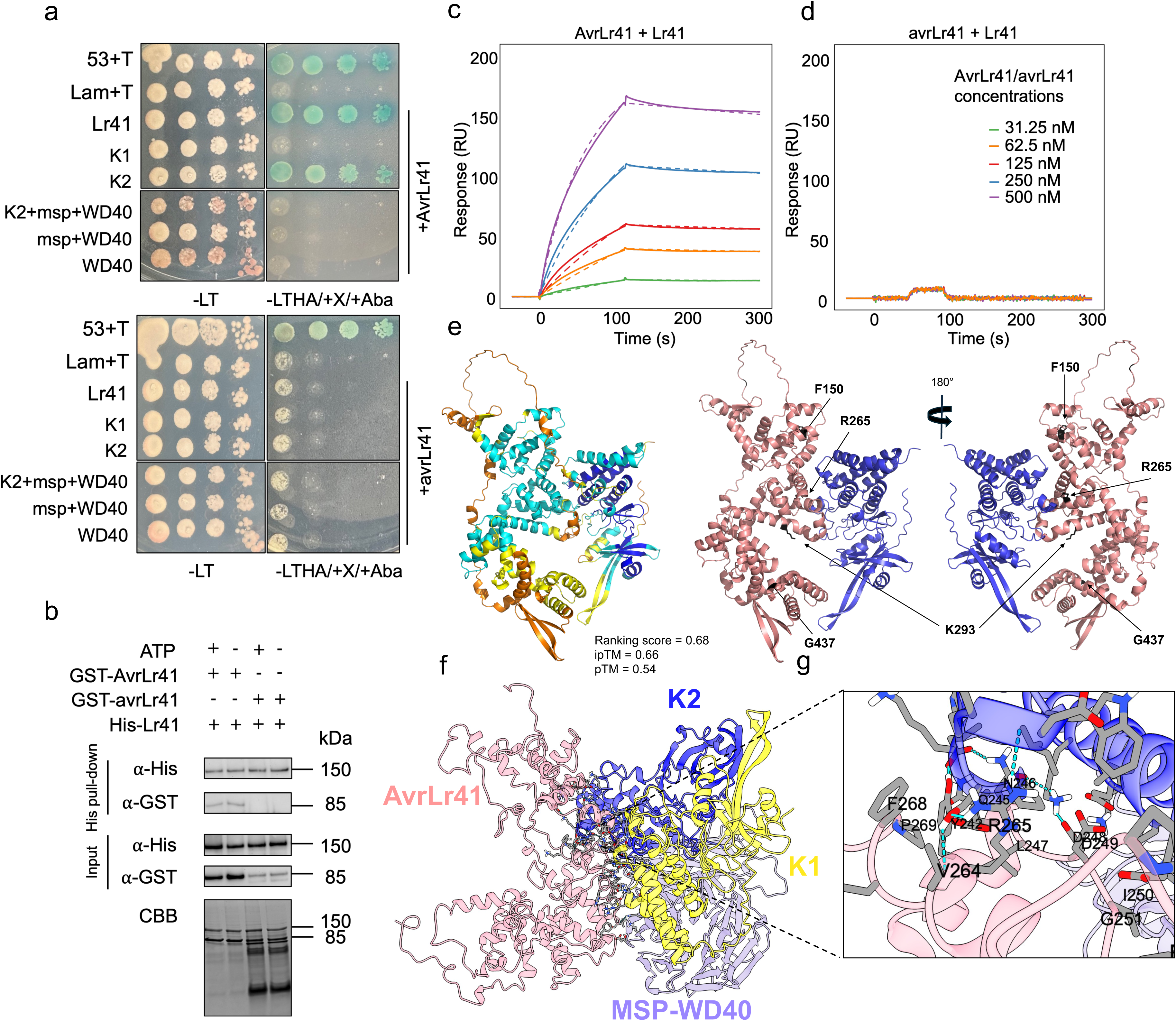
Direct binding of AvrLr41 with Lr41. (**a**) Yeast two-hybrid assay showing direct interaction between AvrLr41 and Lr41. AvrLr41 and avrLr41 were fused to the GAL4 DNA-binding domain (BD), and full-length Lr41 or individual Lr41 domains (K1, K2, K2+MSP, MSP+WD40 and WD40) were fused to the GAL4 activation domain (AD). Diploid yeast strains co-expressing both the AD and BD constructs were selected on synthetic defined (SD) medium lacking leucine and tryptophan (-LT), and interaction-dependent growth was assayed on high stringent SD/-Leu/-Trp/-His/-Ade plates supplemented with X- -gal and Aba. The 53+T pair was served as the positive interaction control, and Lam+T was the negative control. Yeast cultures at an OD_600_ of ∼1.0 were serially diluted ten-fold and spotted on each medium type. Representative photographs were taken after 6 days of incubation. (**b**) *In vitro* His pull-down assay showing that AvrLr41 but not its variant avrLr41 interacts with Lr41 *in vitro*. His-tagged Lr41 and GST-tagged AvrLr41 or avrLr41 were mixed, with ATP added to some samples. CBB: Coomassie Brilliant Blue staining. (**c-d**) Surface plasmon resonance (SPR) analysis of the Lr41-AvrLr41 binding. Sensorgrams obtained by injecting purified GST-AvrLr41 over immobilized His-Lr41 yielded kinetic parameters (ka = 3.07 × 10L ML¹sL¹, kd = 3.21 × 10LL sL¹) corresponding to an equilibrium dissociation constant of ∼10.5 nM. The GST-avrLr41 protein was assessed in parallel. Five concentrations for AvrLr41 and avrLr41 were used as analytes. (**e**) AlpahFold3 (AF3) prediction for the interaction between K2 and AvrLr41. The top-ranked model (ranking score = 0.68; ipTM = 0.66; pTM = 0.53) is shown. Chain-pair ipTM score (0.81 for K2 to AvrLr41) supported a confidently predicted intermolecular interface. The AF3 multimer model in (**e**) is shown with chain-specific coloring to delineate the K2-AvrLr41 interaction. Positions corresponding to naturally occurring AvrLr41 substitutions are labelled. (**f**) Structure model of AvrLr41 (pink) built from cryo-electron microscopy, docking with Lr41 (top-scoring HADDOCK-docked orientation). (**g**) Local view of the docking interface showing the K2 domain positioned along AvrLr41. Cyan dashed lines indicate predicted hydrogen bonds between the docked components. AvrLr41 interface residues are labelled.

To functionally validate *AvrLr41*, we used barley stripe mosaic virus (BSMV)-mediated expression in the Tc and Tc+*Lr41* backgrounds (Fig. 2e). Because of the large size of *AvrLr41*, we designed three partially overlapping fragments spanning the entire gene (Fig. S6) for reconstituting the function of the full-length *AvrLr41* when co-expressed. We observed viral restriction in Tc+*Lr41* leaves when all three fragments were co-expressed (Fig. 2e), whereas the Tc controls developed typical viral symptoms (Fig. 2e). By contrast, four other effector candidate genes from the same GWAS interval failed to trigger an immune response in Tc+*Lr41* when expressed with the BSMV system (Fig. S7). We also overexpressed the candidate *AvrLr41* and its variants in wheat protoplasts using a dual-luciferase defence reporter system^43^. AvrLr41 triggered strong defence reporter activation in Tc+*Lr41* but not in Tc (Fig. 2h), indicating direct recognition of AvrLr41 by the Tc+*Lr41* cells triggering defense responses. The *avrLr41* variant in s692 failed to activate the reporter, comparable to the negative control AvrSr50 (Fig. 2h). As controls, the co-expression of AvrSr50 and Sr50 showed defense induction in both Tc and Tc+*Lr41*, whereas AvrSr50 alone did not (Fig. 2g-h). These results demonstrate that AvrLr41 recognition is *Lr41*-dependent. Notably, AvrLr41 activated defense responses in protoplasts of the synthetic hexaploid L672 carrying *Lr39* (Fig. 2g), reinforcing the notion that *Lr39* and *Lr41* represent the same gene. Together, these results provide direct evidence that AvrLr41 is specifically recognized by Lr41. AvrLr41 thus represents one of the largest fungal Avrs identified to date, comparable in size only to AvrSr35. This finding contrasts sharply with the typically compact effector repertoires of cereal rust fungi^47^.

### Lr41 and AvrLr41 directly interact with each other both *in vitro* and *in vivo*

The *Lr41*-mediated resistance phenotype is an HR, suggesting that direct or indirect recognition of an effector by Lr41 activates defenses. To determine the mode of action of the Lr41-AvrLr41 pair, we first examined it using yeast two-hybrid (Y2H) assays. The tests showed clearly that there was direct interaction between AvrLr41 and Lr41 via binding of the K2 domain (Fig. 3a). The structure combination of MSP-WD40 fusion to K2 (Fig. S1c) prevented K2 interaction with AvrLr41 in yeast (Fig. 3a). When we performed the same assay using the virulent variant avrLr41 from the s692 mutant, there was no interaction with Lr41 in yeast (Fig. 3a). We produced and purified His-Lr41, glutathione S-transferase (GST)-AvrLr41, and GST-avrLr41 (from s692) recombinant proteins (Fig. S10) for *in vitro* pull-down assays; His-Lr41 successfully pulled down GST-AvrLr41, but not avrLr41_s692 (Fig. 3b). We conducted Surface Plasmon Resonance (SPR) to examine the kinetics of interaction between Lr41 and AvrLr41 *in vitro*. We observed direct and high-affinity binding of AvrLr41 to Lr41 when recombinant purified AvrLr41 was applied to immobilized Lr41 (KD ≈ 10.5 nM; ka = 3.07 × 10^4^ M^01^s^01^, kd = 3.21 × 10^04^ s^01^) (Fig. 3c), fitting a 1:1 binding model well (χ² = 4.15). By contrast, the avrLr41 variant failed to bind to Lr41 at all tested concentrations (Fig. 3d), consistent with the lack of interaction between these proteins in yeast (Fig. 3a) and in His pull-down assays (Fig. 3b), and their inability to trigger defense signaling *in planta* (Fig. 2h). Together, these results demonstrate that AvrLr41 directly associates with Lr41 and that this interaction does not occur with the sequence variant avrLr41, supporting a specific recognition of AvrLr41 in *Lr41*-mediated immunity.

### Structural modelling of AvrLr41 recognition by Lr41

Structural predictions of AvrLr41 without the 19-aa signal peptide by AlphaFold 3 (AF3) yielded an overall low-confidence model (ranking score 0.45) (table S6) with three structured helical regions connected by flexible loops (Fig. 3e). The highest ranked complex (at 0.68) involved a K2 and AvrLr41 pairing (Fig. 3e), consistent with the K2-AvrLr41 binding observed in Y2H (Fig. 3c). The two kinase domains of Lr41 share 47% identity, with K1, but not K2, containing conserved kinase motifs^44^ . K1 is thus proposed to be the catalytical component, whereas K2 functions as a pseudokinase domain for effector binding (Fig. 3e).

To gain structural insights into AvrLr41 recognition, we constructed a cryo-electron microscopy (cryo-EM) density map of AvrLr41 at about 7 Å resolution (Fig. S11a-e). The map quality allowed a refined AvrLr41 atomic model (Fig.3e; table S7) built from the experimental density (Fig. S11). The resulting model revealed an extended, multi-domain architecture composed of disordered N-terminal and helical bundles connected by flexible loops (Fig. 3f; Fig. S11b-e), a configuration uncommon among known rust effectors (Fig. S8). The resolved structure is similar to the topology predicted by AF3, despite its low confidence score (Fig. 3e; table S6). Further structural comparison and clustering with ∼23,000 best-modelled proteins from 22 predicted fungal/oomycete secretomes only identified a compact two-member cluster comprising AvrLr41 and a *Pgt* protein with low AF3 prediction confidence (predicted template modeling [pTM]=0.3), with extended connections to additional six low-scoring (bitscore=∼30) proteins from rust and distantly related fungi (Fig. S8). This finding indicates that AvrLr41 only marginally resembles the structures of other fungal effector proteins. We next examined how AvrLr41 directly binds to Lr41. We used the predicted full-length Lr41 structure (pTM=0.6; table S6; Fig. 3f) to dock against the AvrLr41 model (Fig. 3f). The best-fit complex (HADDOCK scores, table S8) positioned K2 as the primary interface, forming an extensive contact surface with AvrLr41 (Fig. 3f-g). By contrast, the K1 and C-terminal MSP-WD40 domains were oriented distally, suggesting a K2-mediated effector recognition. AvrLr41 appears to adopt a complex fold that is distinct from the compact architecture of most rust Avrs^45^ and directly interacts with the Lr41 K2 domain.

### An NLR helper is required for Lr41-AvrLr41 mediated immunity

The resistance mediated by Lr41 is intriguing because it involves direct binding with such an unusual Avr. Current models of plant immunity describe sensor-helper systems in which resistance often depends on helper NLRs found at proximity to their cognate sensor genes^9,28,29^. Despite the direct interaction between Lr41 and its paired effector AvrLr41, no immediate HR was observed when Lr41 and AvrLr41 were co-expressed in *Nicotiana benthamiana* (Fig. 4; Fig. S14a-b and f; Fig. S15), suggesting that additional host factors are required to execute *Lr41*-mediated resistance. To identify potential candidate helpers, we generated a contig-level genome assembly of Tc+*Lr41* and inspected the ∼1 Mb region surrounding *Lr41* (Fig. 4a). Notably, one gene located 616 kb upstream of *Lr41* (Fig. 4a) shows sequence similarity to NLRs but encodes only a truncated form. Its transcript was confirmed by RNA-seq (Fig. S12a) and cDNA amplification followed by sequencing. Surprisingly, this contrasts with an intact NLR ortholog present at the corresponding locus in *Ae. tauschii* TA1675, which is also expressed (Fig. 2b; Fig. S12a). Thirteen SNPs distinguish the two orthologs (Fig. 2b), indicating that additional recombination or selection events occurred either prior to or following the introgression of the *Ae. tauschii* 2DS segment conferring *Lr41* resistance in Tc+*Lr41*. The truncation results from a single-base deletion at position +239 in Tc+*Lr41* (Fig. 2b; Fig. S12a-b), introducing a premature stop codon at 561 bp immediately upstream of the NB-ARC region. This produces a truncated protein consisting of two N-terminal coiled-coil (CC) domains followed by an intrinsically disordered C-terminal extension (Fig. S12 a-d). Beyond the *Ae. tauschii* orthologs, the closest related sequence is a protein with truncated CC domains from *Roegneria kamoji* (60% identity, 12% gaps; Fig. S17a-b): these CC domains share homology with those of *Thinopyrum intermedium* and the barley Rph1 (Fig. S17a-b). Phylogenetic analysis placed these CC sequences in a distinct clade separated from all major N-terminal lineages of both canonical and atypical NLR families (Fig. S17a-b).

**Fig. 4.**
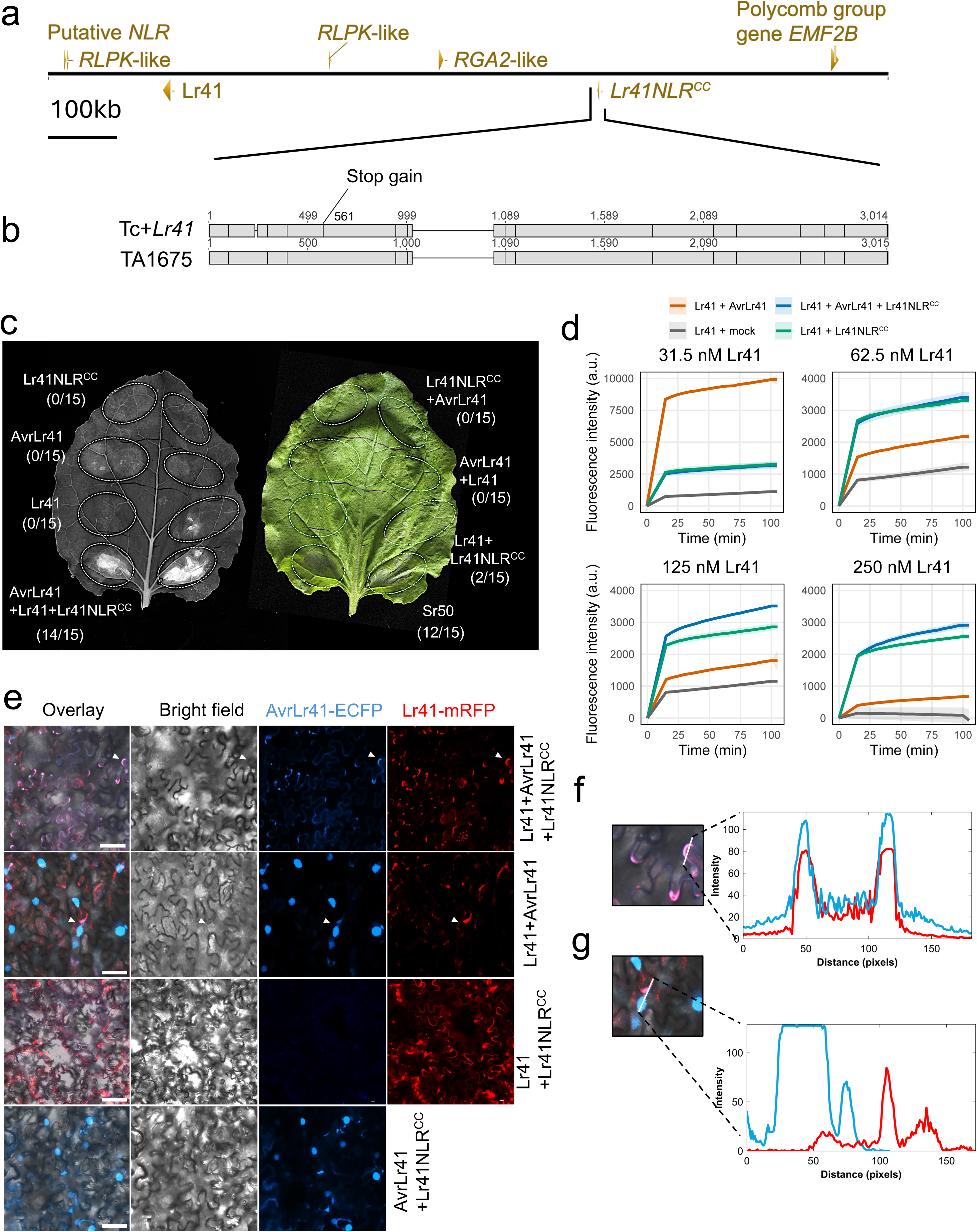
**A truncated helper NLR links AvrLr41 recognition to Lr41-mediated immune signaling.** (**a**) Diagram of the genomic position of the *Lr41* locus and neighbouring genes in Tc+*Lr41* and TA1675. RLPK: Receptor-Like Protein Kinase; RGA2: Resistance Gene Analog 2; EMF2B: Embryogenic Flower2B (**b**) A truncated NLR (Lr41NLR^CC^) encoding gene resides ∼600 kb upstream of *Lr41* and shares synteny with a complete NLR ortholog in TA1675. Gene models of *Lr41NLR^CC^* in Tc+*Lr41* and its ortholog in TA1675 show a single-base deletion at position +239 that introduces a stop codon, ending at +560. Vertical lines denote SNPs. (**c**) Co-expression assays in *Nicotiana benthamiana* demonstrating that the *Lr41*-mediated hypersensitive response requires both AvrLr41 and Lr41NLR^CC^. Sr50 served as an autoactivation positive control. The numbers in parentheses indicate the number of replicates showing cell death out of the total replicates. (**d**) *In vitro* kinase assays showing Lr41 catalytic activity toward AvrLr41 and its modulation by Lr41NLR^CC^. Time-resolved fluorescence measurements at different Lr41 concentrations show the phosphorylation of AvrLr41 by Lr41. Lr41 alone served as background control. Fluorescence intensity is shown as arbitrary units (a.u.) after normalization by background subtraction. (**e**) Translocation of AvrLr41 requires the Lr41-Lr41NLR^CC^ sensor-helper module. Confocal microscopy of *N. benthamiana* epidermal cells expressing *35S:AvrLr41-ECFP* and *35S:Lr41-mRFP*, with or without *Lr41NLR^CC^*. A Nikon inverse confocal microscope was used with ECFP excitation/emission at 433/475 nm and mRFP with excitation/emission at 584/607 nm. The white arrowheads point to the regions used for Lr41-AvrLr41 colocalization analysis. (**f-g**) Analysis of Lr41-AvrLr41 co-localization. Fluorescence-intensity profiles were extracted along the indicated scan paths for the regions indicated by arrows, showing colocalization of Lr41 and AvrLr41 (**f**) and lack of nuclear colocalization (**g**).

To assess the function of the truncated NLR, we silenced *Lr41NLR^CC^* using BSMV-mediated VIGS in Tc+*Lr41* plants (Fig. S12e-f). Silencing reduced *Lr41*-mediated resistance to *P. triticina* isolate PBJJG accompanied by greater uredinial development (Fig. S12e) and lower *Lr41NLR^CC^*expression (Fig. S12f), indicating that *Lr41NLR^CC^* contributes to the *Lr41*-mediated resistance. To explore whether the expressed NLRs at the *Lr41/39* locus contribute to Lr41-AvrLr41-mediated defense, we co-expressed *Lr41*, *AvrLr41*, and different NLR variants in *N. benthamiana* (Fig. 4c; Fig. S13; Fig. S14a-b; Fig. S15) and barley protoplasts (Fig. S14f). Both *Lr41NLR^CC^* (Fig. 4c; Fig. S14b) and its *Ae*.

*tauschii* ortholog *AetNLR* (Fig. S14a-b) elicited a HR only when co-expressed with *Lr41* and *AvrLr41* in *N. benthamiana*, comparable to the positive autoactivation control *Sr50* (Fig. 4c; Fig. S14a-b; Fig. S15). A similar HR pattern could also be observed in barley protoplast transiently expressing *Lr41NLR^CC^* or *AetNLR* with both *Lr41* and *AvrLr41* (Fig. S14f). No other combinations of *AetNLR^CC^*, *Lr41*, *AvrLr41*, *Lr41NLR^NBS^*/*AetNLR^NBS^*, or any gene alone could trigger HR (Fig. 4c; Fig. S14a-b; Fig. S15). As a negative control, we included the leaf rust resistance NLR gene *Lr13* from wheat^46^. Co-expression of *Lr41*, *AvrLr41* and *Lr13* did not elicit any HR in *N. benthamiana* (Fig. S15), indicating that helper activity is specific. Lr41NLR^CC^ interacted with Lr41 and its MSP-WD40 domain but not with AvrLr41 in yeast, whereas AetNLR only bound directly to the MSP-WD40 domain of Lr41 (Fig. S16). The MSP-WD40 domain showed broad interactions with multiple NLR variants (Fig. S16). These interaction patterns are consistent with the HR elicited by both *Lr41NLR^CC^* and *AetNLR*, and support MSP-WD40 as a primary contact site for the helpers.

### A truncated helper NLR couples AvrLr41 recognition to Lr41 localization and kinase activity

CC-NBS-LRRs have been shown to form oligomeric complexes at the plasma membrane, yet it is currently unknown where and how TPKs engage in signaling with helper NLRs^28,29^. One intriguing finding was that when *Lr41NLR^CC^* was co-expressed with *Lr41* and *AvrLr41*, AvrLr41 relocated from the nucleus (Fig. 4e and g) to plasma membrane-associated and cortical cytoplasmic regions (Fig. 4e-f), where it co-localized with Lr41 at the cell borders, stomata, and chloroplast periphery (Fig. 4e-f). Distinct AvrLr41-ECFP puncta were also observed along the plasma membrane (Fig. 4e; Fig. S14e). AvrLr41 contains predicted nuclear localization signals (Fig. S9b-c), consistent with its nuclear localization. Fluorescence intensity analyses confirmed spatial overlap of Lr41-mRFP and AvrLr41-ECFP signals only when all three proteins were present (Fig. 4e). In contrast, AvrLr41 remained in the nucleus when co-expressed with either Lr41 or Lr41NLR^CC^ alone (Fig. 4e). These results indicate that Lr41, AvrLr41, and Lr41NLR^CC^ may act cooperatively, in line with the physical interactions observed both *in vivo* and *in vitro* and the HR observed upon co-expression of all three. Consistently, AetNLR displayed a similar function to work together with Lr41 to facilitate translocation of AvrLr41 from the nucleus to plasma membrane-associated and cortical cytoplasmic regions and stomata (Fig. S14c-d). Interestingly, in the presence of all three recombinant proteins *in vitro*, rather than with only Lr41 and AvrLr41, a pore-like structure was clearly observed via cryo-EM (Fig. S18a-h). How the pore-shaped particles assemble and whether they represent resistosome-like oligomers (Fig. S18a-f) will be determined in future studies.

To further examine whether the truncated helper NLR influences Lr41 signaling output, we assessed Lr41 kinase activity *in vitro* in the presence of AvrLr41 and/or Lr41NLR^CC^. We quantified *in vitro* Lr41 kinase activity across a range of Lr41 concentrations while maintaining AvrLr41 at a fixed level and, in separate reactions, including Lr41NLR^CC^. Lr41 exhibited substantially higher kinase activity in the presence of AvrLr41 and/or Lr41NLR^CC^ than it did on its own (Fig. 4d), consistent with a role in the transphosphorylation of other proteins. At the lowest Lr41 concentration, the Lr41-AvrLr41 complex exhibited substantially higher activity than either Lr41-Lr41NLR^CC^ or the ternary Lr41-Lr41NLR^CC^-AvrLr41 mixture (Fig. 4d). When Lr41 concentration increased to 250nM, kinase activity decreased across all reactions, but to varying degrees. The reduction was most pronounced for Lr41-AvrLr41, whereas reactions containing Lr41NLR^CC^ showed a more gradual decline (Fig. 4d). Notably, inclusion of Lr41NLR^CC^ preserved a moderate level of kinase activity across all Lr41 concentrations and resulted in stronger signals than Lr41-AvrLr41 alone at higher Lr41 concentrations, suggesting that the presence of AvrLr41 no longer alters the catalytic activity of Lr41 once Lr41NLR^CC^ binds (Fig. 4d). This equivalence further suggests that Lr41NLR^CC^ may modulate Lr41 kinase activity towards AvrLr41, where Lr41NLR^CC^ binding induces a conformational change in Lr41 that allosterically uncouples the effector-binding domain from catalytic activation. In this scenario, AvrLr41 may still bind the K2 domain, as shown in yeast (Fig. 2c), but this binding no longer stimulates kinase activity. These results, together with the ability of Lr41NLR^CC^ to reconstitute HR with Lr41 and AvrLr41, and the coordinated relocalization of AvrLr41 *in planta*, support a model in which Lr41-mediated immunity may require an auxiliary helper component to translate effector recognition into downstream defense activation at the plasma membrane-associated locations.

## Discussion

Pathogen effector perception in plants is often framed by NLR-centered models, in which Avr recognition is direct or indirect via guard/decoy mechanisms, and culminates in resistosome assembly and cell death^1–10^. This study extends these models by identifying an unconventional R-Avr pair, namely Lr41-AvrLr41, underlying wheat leaf rust resistance. First, we characterized Lr41 as a TKP-type R protein with C-terminal fusion domains. Second, we identified its paired AvrLr41 as a structurally highly divergent Avr. Third, we uncovered a non-canonical signaling module in which Lr41 detects AvrLr41 through its pseudokinase domain and signals via alternative helpers, including an unusually truncated NLR. In our current model, the pseudokinase domain of Lr41 acts as the sensor and the C-terminal fusion as a scaffold to recruit the helper(s) for coordinated detection of AvrLr41 (Fig. S19). Our findings extend the emerging paradigm that TKPs operate as Avr sensors that enable immunity execution with partnered NLRs^28,29,47^. Unlike Sr62 and WTK3, which lack additional fusion domains and bind effectors via their kinase domains, Lr41 relies on pseudokinase-mediated recognition and likely employs its C-terminal domain to engage either a minimal CC-only helper or a canonical full-length NLR. The apparent catalytic activity of the K1 domain of Lr41 suggests a possible phosphorylation step coupled with K1-mediated effector sensing and helper recruitment. The coexistence of a minimal CC-only helper and a canonical NLR reveals unexpected plasticity and evolutionary modularity in TKP-NLR signaling assemblies.

Remarkably, the truncated helper Lr41NLR^CC^, lacking the NB-ARC and LRR regions and even the third and fourth α-helices of a standard CC domain, remained competent to facilitate Lr41-dependent HR in *N. benthamiana* and barley, while the full-length *Ae. tauschii* ortholog AetNLR can also function as a helper. The ability of an NLR derivative of two α-helix bundle CC to substitute for canonical helpers defines a previously unrecognized flexibility in plant immune architecture. Both helpers could interact with the Lr41 MSP-WD40 domain and enable AvrLr41 translocation to plasma membrane-proximal sites, indicating assembly of a membrane-associated protein complex. Such kinase-type R-Avr recognition model diverges from the standard NLR activation yet aligns with the membrane-proximal resistosome paradigm exemplified by NLR oligomerizations^27^. Although pore-like particles were observed *in vitro* when all three recombinant proteins were present (Fig. S18), whether these particles represent functional resistosome-like oligomers remains to be determined. The CC bundle is known to execute immune signaling once relieved from autoinhibition. Unlike the CC domains from canonical CNLs (CC-NLRs) such as ZAR1^48^, MLA10^49^, Sr33 and Sr50^50^, which independently trigger HR, the CC domains associated with Lr41 appeared to function primarily in complex assembly, not autonomous cell death induction.

This study resolves the long-standing ambiguity regarding whether *Lr39* and *Lr41* are allelic or identical ^40–42^; however, their breeding histories appear to have diverged. Compared to the *Ae. tauschii* donor line TA1675 for *Lr39*, Tc+*Lr41* showed substantial variations over the *Lr41* interval, most notably with the truncated CC-only helper, whereas *Ae. tauschii* retains an intact ortholog. Historically, *Lr39* and *Lr41* were introgressed from distinct *Ae. tauschii* accessions, yet both confer identical specificity and infection type against diverse *P. triticina* pathotypes differing in virulence for *Lr41*^40,51^. Evidence for *Lr39* in *Ae. cylindrica* suggests an ancient origin predating the emergence of this species or interspecific exchange within the D-genome lineage between *Ae. tauschii* and *Ae. cylindrica*^41^. Despite these evolutionary differences, resistance remains fully functional, with both Lr41NLR^CC^ and AetNLR capable of partnering with *Lr39/Lr41*. Furthermore, silencing AetNLR alone does not abolish *Lr39* resistance^22^, consistent with functional redundancy among helper NLRs. If there are additional helpers in *Ae. tauschii*, loss or lower expression of a single helper gene would not be sufficient to disrupt *Lr41*/*Lr39*-mediated immunity. Future studies overexpressing different NLR helper variants together with *Lr41*, along with finding other potential helpers, will be required to fully elucidate *Lr41*-dependent defense, as proposed in models where expanded NLR network partners provide flexible immune signaling to counteract rapidly evolving plant pathogens^2,52^. The historical and genomic context refines both the mechanistic interpretation of sensor TKP with interchangeable helpers and the translational path toward durable stacking of non-redundant resistances alongside redundant gene contents.

## Methods details

### Plant and fungal materials

The bread wheat cultivar ‘Tc’ and the NIL Tc+*Lr41* are maintained at the Plant Breeding Institute (PBI), Cobbitty, The University of Sydney. Wild-type and EMS mutant plants were grown in greenhouse at 16-20 °C; when they reached the two-leaf stage, seedlings were inoculated with the leaf rust pathotype 64-11 (University of Sydney culture no. 473; s473) (table S5). Infection types were recorded 10-14 days post inoculation (dpi). Isolate s473 and other *P. triticina* isolates were sourced from liquid nitrogen storage at the PBI Cobbitty, The University of Sydney. For VIGS, the US pathotype PBJJG, maintained at Montana State University, USA, was used. Leaf rust virulence and avirulence phenotypes are listed in table S5. *N. benthamiana* plants were grown under a 16-h light/8-h dark photoperiod with 70% humidity at 25 °C. For protoplast isolation, the wheat cultivar Tc, the NIL Tc+*Lr41,* and the synthetic line L576 were grown in controlled environmental chambers at 21 °C under a 16-h light/8-h dark photoperiod, with light intensity maintained at 100-150 µmol m^−2^ s^−1^ at canopy level and the relative humidity set to 80%.

### Mutagenesis

Ethyl methanesulfonate (EMS) mutagenesis was conducted on 2,100 seeds of the NIL Tc+*Lr41* treated with 0.5% or 0.6% (v/v) EMS (Sigma-Aldrich, M0880) as previously described^46^. In brief, seeds in batches were wrapped in cheesecloth and soaked in 700 mL of EMS solution for ∼18 h on a shaker, followed by extensive washing under running water for 2 h before being sown in pots. M2 seedlings from up to six single spikes per M1 plant were inoculated with pathotype 64-11; candidate mutants in *Lr41* were identified as susceptible seedlings in segregating single spike progenies. Susceptible M3 *lr41* mutant plants and their resistant siblings were further progeny-tested to generate M4 plants. Thirty-nine independent *lr41* mutants from M4 plants, which were confirmed by progeny tests, were selected for PCR testing and DArTseq (Diversity Arrays Technology Pty Ltd, Australia). Six M3 or M4 mutants were used for MutRenSeq (Arbor Biosciences, MI). An additional three mutants, i.e. M3.93, M4.38, and M4.26, were used for amplicon amplification and Sanger and Nanopore sequencing.

### High molecular weight DNA extraction and sequencing

Fresh tissue from the third and fourth leaves from four-week-old wheat plants was flash-frozen in liquid nitrogen and ground to a fine powder. Approximately 20 mL of CTAB buffer containing 1% (v/v) β-mercaptoethanol, 200 µL of 20 mg/mL RNase A, and 200 µL of 20 mg/mL Proteinase K were added to each sample. The samples were incubated at 65 °C for 1 h with gentle inversion every 10 min. Extracts were cooled on ice, mixed with an equal volume of chloroform:isoamyl alcohol (24:1, v/v), and rotated at 60 rpm for 10 min before centrifugation at 6,000 × g for 20 min at 10 °C. Seventeen milliliters of the supernatant were recovered and subjected to a second chloroform:isoamyl alcohol extraction. Fifteen milliliters of supernatant was then combined with two volumes of cold isopropanol and 0.1 volume of 3 M sodium acetate. Samples were gently inverted and incubated at room temperature for 5 min to allow DNA to precipitate. Genomic DNA was pelleted by centrifugation at 10,000 × g for 5 min and resuspended in 10 mM Tris buffer (Tris-HCl, pH=8). Following another round of isopropanol and chloroform purification, DNA pellets were washed with 80% (v/v) ethanol, centrifuged again at 10,000 × g for 5 min, air-dried, and resuspended in 400 µL Tris buffer. For PacBio HiFi sequencing of Tc+*Lr41*, the high molecular weight DNA sample was sequenced on a PacBio Revio System sequencer using two Revio SMRT Cells, which generated 183.1 Gb HiFi reads (Novogene, Singapore). For genome assembly, hifiasm (v0.19) with default parameters was used. Assembly completeness was assessed using the embryophyta_odb12 database with Compleasm (v0.2.7)^53^, which identified 2,026 BUSCO groups, of which 98.28% were complete (8.05% single-copy, 90.23% duplicated), 0.89% were fragmented, and 0.84% were missing.

For MutRenSeq, six independent susceptible M3 or M4 mutants (285M4.12, 348M4.14, 2419M3.30, 1392M3.18, 1399M3.23, 202M4.4) were selected. The second leaves from seedlings of each mutant and Tc+*Lr41* were collected for genomic DNA at 10 dpi. NLR target enrichment using the *Triticeae* RenSeq bait library, fragment capture, and library preparation were performed by Arbor Biosciences (MI, USA), and sequenced on an Illumina NovaSeq 6000 S4 instrument. For each mutant, 6.8, 9.6, 52.7, 69.8 and 33.9 M 150-bp paired-end (PE150) clean reads were produced, respectively. An additional mutant (1392M3.18) and the NIL Tc+*Lr41* were processed for PacBio HiFi (CCS) sequencing following bait capture. A total of 2.1 M and 2.3 M demultiplexed CCS reads with a median length of 3.6 kb and 3.9 kb for the mutant and Tc+*Lr41*, respectively, were produced from two HiFi 8M Sequel II cells (PacBio). The MutRenSeq pipeline was followed to identify candidate NLRs. In brief, the pipeline extracts EMS-type G/C-to-A/T transitions in the sequence of the EMS mutants relative to the *Ae. tauschii* TA1675 genome. The TA1675 reference contigs were aligned to the source sequences of the bait library (https://github.com/steuernb/MutantHunter/blob/master/Triticea_RenSeq_Baits_V3.fasta.gz) using BLASTn. Aligned contigs were then used as reference for mapping the reads from the mutants, as well as those of the WT as a positive control. Short-reads and long-reads were mapped to the TA1675 genome assembly using bwa (v0.7.18) and minimap2 (v2.24) -ax map-hifi. The alignment results were further processed into BAM format using SAMtools (version 1.9) with default parameters. The mapping data and annotation of NLR type were combined to determine candidate contigs based on EMS-type mutations in the same contig.

### RNA extraction and sequencing analysis

For mRNA-seq, total RNA was extracted using the RNeasy Plant Mini Kit (Qiagen, Germany) from pooled seedlings for Tc and Tc+*Lr41*, respectively, harvested at 0, 6, 24, 48, 72, 96 and 168 hours post inoculation (hpi) with an *Lr41* avirulent pathotype (s473). mRNA samples were enriched using oligo(dT) beads (AGRF, Australia), and sequencing was conducted with Illumina NovaSeq 6000 instrument (AGRF), producing an average of 200 M reads per replicate for Tc and Tc+*Lr41*. Raw reads were mapped to the TA1675 reference genome with hisat2 (default settings). This analysis revealed an average of ∼1.7 times more uniquely mapping reads for Tc+*Lr41* (∼ 8.5%) than for Tc (∼4.8%), and an overall alignment rate of ∼12 % (Tc+*Lr41*) or 8% (Tc). For expression analysis to identify putatively expressed candidate *Lr41* as well as NLR-encoding genes, DEseq2^54^ was used with IWGSC CS RefSeq v2.1 (CSv2.1) as the reference genome. To enable transcript investigation, strand-specific mRNA-seq data from the TA1675 line^22^ were used. Raw reads were trimmed with fastp (v0.23) and mapped to the TA1675 primary genome assembly using Hisat2 with the strand-specific option set as FR. About 55 M (over 91%) of all reads were concordantly mapped with read alignment rates over 95%.

### Comparative genomics

An “intersection-of-expression” workflow was developed to identify *Lr41*: screening all potential variations in the TA1675 2DS region by comparing *T. aestivum* and *Ae. tauschii* lines with or without *Lr41*. mRNA-seq data generated from the NIL Tc+*Lr41*, its recurrent line Tc, and *Ae. tauschii* were analyzed. Sequence variations in the putative NLR genes in the TA1675 2DS region were explored. The TA1675 2DS genomic region was annotated by BLAST searches against a plant NLR database^55^. All putative NLR sequences were used as reference for the alignment of RNA-seq reads. To determine the interval on 2DS harboring *Lr41*, direct genome-genome alignments were performed across the contig-level assembly of TA1675, chromosome 2D of CSv2.1, and the *Ae. tauschii* AL8/78 (Aet5.0) genome assembly using minimap2. Because the sequence divergence may be high particularly over the 2DS region, the option -asm20 was used. The closest linked marker *Xbarc124* and two other linked markers^42^ were anchored to the above references. The distal end regions for the chromosome segment 2DS from all reference assemblies were extracted using SAMtools and realigned with minimap2 with the -asm20 option. For exploring the synteny of the distal ends of the 2DS segment between TA1675 and CS, the paf alignment output was plotted with the pafr package (v0.02). Combined bulk RNA-seq reads were aligned to the TA1675 reference genome with Hisat2, and transcript structures were assembled using StringTie (v2.2.1) with default parameters. GTFs files were merged into a unified annotation, and transcript abundance was quantified with Cufflinks and normalized with Cuffdiff^56^. Expression matrices were clustered by Euclidean distance with Ward’s method, and heatmaps were generated with the heatmap.2 function of the R package gplots. In addition, normalized read depth across the distal 2DS interval for all transcript coordinates was calculated with SAMtools and visualized in R (v4.3.2) using ggplot2 as line and area plots with locally estimated scatterplot smoothing. Freebayes with vcffilter (QUAL > 20) and vcftools (vcf-annotate -f | vcf-contrast | vcf-query -f) were used to extract homozygous SNPs and their depth across the transcript regions.

### Rust pathogenomics

For mapping the sequences from leaf rust isolates, a revised fully-phased reference genome for s473 was used^57^. This method allowed a fast examination of pseudo-allelic candidates from the two haplotype-resolved genomes of the mutants where alleles (if any) should encode a nonfunctional variant (avrLr41) of the wild-type effector (AvrLr41). Reads were mapped to the two haplotype-resolved genomes separately using BWA. The resulting outputs were processed with SAMtools and visualized with the Integrative Genomics Viewer (IGVv2.12). The haplotype variant caller freebayes (v1.3.2) was then used to generate variant outputs in Variant Call Format (VCF) containing SNP and small insertion/deletion (InDel) information. The outputs were filtered using the vcffilter tool in vcflib (v1.0) with parameters DP>20 QUAL>20 to only retain high-quality mapping. Vcffilter (--maf0.02 --miq10 -no-indels) was used to filter VCF files as the input for GWAS analysis. For haplotype GWAS, a linear mixed model and Wald test with 52 *P. triticina* RNA-seq and genome-seq datasets were used. PLINK and bcftools with linear models in GEMMA were used, and frequentist Wald test, likelihood ratio and score were performed in vcf2gwas^58^. A total of 8,041 and 8,517 SNPs (MAF ≥ 0.02) from haplotype 1 and haplotype 2, respectively, were analysed.

### Phylogenetic analysis

The sequences of 370 kinase domain-containing proteins belonging to the PANTHER subfamily PTHR45707, mainly from wheat, barley, and *Brachypodium distachyon*, were retrieved and aligned with mafft. Phylogenetic analysis was performed with qitree2 (v2.0) using the aligned protein sequences as input and the maximum-likelihood method. The best-fit amino acid substitution model was selected with ModelFinder (-m MFP). Branch support was assessed with 1,000 ultrafast bootstrap replicates (-bb 1000) and 1,000 SH-aLRT tests (-alrt 1000).

### Structural similarity prediction, molecular networking, and molecular docking

For predictions of protein structure, AlphaFold 3 was installed locally using docker and 5-6 seeds were run. For structural molecular networking, structural homologs were identified using FoldSeek (v.2024-04) against AlphaFold2-predicted secretomes from 22 fungal and oomycete pathogens, including *Pgt* and *Pst*^45,59^. Pairwise structure-structure alignments were performed using FoldSeek easy-search with default settings, producing ∼6.2 million alignments with the bit scores used to quantify fold similarity. These results were filtered to retain matches with bit score ≥ 30 for meaningful structural similarity. The resulting filtered edge list (query, target, bitscore) was clustered using the mcl package (v22-282)^60^. Edge weights were transformed as −log_10_(bit score) during clustering to enable stochastic matrix scaling (mcxload --stream-mirror --stream-neg-log10), followed by mcl -I 2.0 for inflation-based partitioning of structural communities. For visualization, raw bit scores were used to scale edge color and thickness, preserving direct comparability to FoldSeek outputs. Networks were plotted with NetworkX and Matplotlib in Python. Protein-protein molecular docking was performed using HADDOCK v2.4 (https://wenmr.science.uu.nl/haddock2.4/) with default parameters. The interaction interface was defined using Ambiguous Interaction Restraints (AIR). For AvrLr41, the cryo-EM model was used and residues at positions 260-272 (signal peptide removed) were pre-selected as interface drivers. For Lr41, the AlphaFold3 model was used where active residues were selected based on the AIR specification generated for this run and included amino acid (aa) positions 539-651 of the K2 domain, with passive residues automatically assigned by HADDOCK. Docking was carried out through a standard three-stage protocol: rigid-body minimization, semi-flexible simulated annealing, and final refinement in explicit solvent. The resulting models were clustered using the Fraction of Common Contacts method with a 0.6 cutoff and a minimum cluster size of four structures. Clusters were ranked according to the HADDOCK score, calculated as a weighted combination of van der Waals, electrostatic, desolvation, and restraint energies, averaged over the best-scoring structures within each cluster. The highest-ranking cluster (HADDOCK score = −63.807±19; table S8) was considered the most favorable docking solution based on both energetic and structural consistency. Models were visualized in ChimeraX (v1.10).

### Molecular cloning

The full-length coding sequences of *Lr41*, *AvrLr41*, and its mutant *avrLr41* (s692) were synthesized with 20-bp flanking ends homologous to the 5L and 3L *Bam*HL/*Hind*L digestion sites in the pGADT7 and pGBKT7 vectors (Clontech, Takara Bio, Japan) and assembled via Gibson assembly in the *Saccharomyces cerevisiae* strains Y187 and Y2HGold. The *Lr41* full-length coding sequence was split into two fragments with a 30-bp overhang at the break site (table S9). The coding sequences of *Lr41* truncated fragments, including those encoding Kinase domain 1 (aa 1-349), Kinase domain 2 (aa 351-670), msp-WD40 fusion (a 671-1,131), the WD40 repeats (aa 819-1,131), both kinase domains (aa 1-670), and K2-msp-WD40 (aa 350-1,131), were synthesized and assembled into the pGADT7 vector by Gibson assembly using the yeast strain Y187. Coding sequences of the truncated NLR^CC^ (561bp) and NLR^NB-ARC-LRR^ (2,457 bp) from Tc+*Lr41* cDNA were PCR amplified using Phusion DNA Polymerase (Thermo Fisher Scientific, Australia) with primers listed in table S9, and cloned into both the pGADT7 and pGBKT7 vectors by Gibson assembly. The full-length coding sequence of the homolog of the NLR candidate in TA1675 (*AetNLR*) with an *Lr41*-proximal locus similar to *Lr41NLR^CC/NB-ARC-LRR^* was also synthesized and assembled into the pGADT7 and pGBKT7 vectors. To test interaction between N-terminal and C-terminal truncations of the AetNLR candidate with Lr41 or AvrLr41, the sequences encoding AetNLR CC and NB-ARC-LRR (AetNLR^CC^ and AetNLR^NB-ARC-LRR^) were PCR-amplified using primers (table S9) with the same overhangs as mentioned above, and assembled into the pGADT7 and pGBKT7 vectors. For Gibson assembly, yeast transformation was performed using 40% (w/v) polyethylene glycol (PEG) 4000 and 1 M LiAc; selection for auxotrophic growth was performed on synthetic defined (SD) medium lacking either leucine or tryptophan.

Positive transformants were confirmed by PCR using Phire Plant Direct PCR Master Mix (Thermo Fisher) and vector-specific primers (table S9). All assembled plasmids were extracted from positive colonies, retransformed and extracted from TOP10 *E.* coli using the Qiagen Mini Plasmid Extraction Kit, and confirmed with whole plasmid sequencing (AGRF). For subcellular localization and *in planta* transient expression, the full-length coding sequences without the stop codon of *Lr41*, *AvrLr41*, the Tc+*Lr41* version of *NLR^CC^* and *AetNLR* along with its truncations *AetNLR^CC^* and *AetNLR^NB-ARC-LRR^*were PCR-amplified from yeast expression vectors and inserted into pDONR and pTOPO-D-Entry vectors, followed by LR reaction and insertion into binary vectors under the control of the CaMV 35S promoter, yielding the constructs *35S:Lr41-mRFP*, *35S:AvrLr41-YFP*, and 35*S:NLR-3xHA*. A similar strategy was employed to generate *35S:Sr50-3xHA* for use as a control together with pGFPGUSplus. For *in vitro* protein production, the full-length coding sequences of relevant genes were synthesized (GenScript, Singapore) and inserted into the cloning vectors pGEX-6P-1 and pET30a to generate the tagged fusion proteins GST-AvrLr41, GST-avrLr41, His-Lr41, and SUMO-NLR^CC^. For wheat protoplast cell death assays, Golden Gate assembly was used. In brief, gBlock fragments of *AvrLr41* and its nonfunctional mutant allele (*avrLr41*_s692) with *Bsa*I overhangs at both ends were cloned into the pWDI expression vector using *Bsa*I-based Golden Gate assembly (25 cycles of 37 °C digestion, 16 °C ligation, followed by a final incubation at 50°C for 5 min and 80°C for 5 min to inactivate *Bsa*I). Reaction products were introduced into *E. coli* by heat shock. All colonies were screened by colony PCR, and positive clones were validated by restriction digestion and confirmed by whole-plasmid sequencing on an ONT MinION (Oxford Nanopore).

### Recombinant protein production

Recombinant Lr41, AvrLr41, and Lr41NLR^CC^ were produced using codon-optimized fragments as aforementioned in molecular cloning. The coding sequence of the *AvrLr41* nonfunctional variant *avrLr41_s692* was not codon optimized. The full-length *Lr41* coding sequence was cloned into pET30a using *Nde*I/*Hind*III to add an N-terminal 6×His tag followed by a TEV protease recognition site upstream of Lr41. The resulting construct was introduced into BL21 Star (DE3) (Thermo Fisher Scientific) and protein production was carried out at 15 °C for 16 h. To increase solubility and avoid aggregation, 10-L culture scale-up was used for cell cultures induced with 0.5 mM IPTG and grown at 15 °C for 16 h. Cell pellets were resuspended in 2 L lysis buffer (50 mM Tris-HCl, 500 mM NaCl, 1% [v/v] Triton X-100, 1 mM DTT, 10% [v/v] glycerol, pH 8.0), lysed by sonication, and clarified by centrifugation. Lr41 was purified by Ni²L-NTA chromatography, eluting predominantly in the 500 mM imidazole fraction, followed by Superdex 200 size-exclusion chromatography (SEC) (Cytiva). HPLC was used to estimate the proportion of monomer in the purified fraction prior to scale-up. Two rounds of SEC were performed. To stabilize the protein during upstream purification, buffers initially containing 10% (v/v) glycerol and 100 mM arginine were included during dialysis to enhance monomer recovery. The final Lr41 preparations were buffer-exchanged against a glycerol-free buffer

(50 mM Tris-HCl, 150 mM NaCl, pH 8.0) and reached a purity of 85%. AvrLr41 and avrLr41_s692 were produced as N-terminal GST-TEV fusions in DE3 and Rosetta *E. coli* cells, respectively. Cells with each construct were induced with 0.5 mM IPTG and incubated at 15 °C for 16 h in 1 L of LB. Cells were lysed by sonication, and the fusion proteins were purified by glutathione affinity chromatography (GeneScript, L00206) and eluted in 50 mM Tris-HCl, 150 mM NaCl, pH 8.0. SDS-PAGE and immunoblot analysis showed that GST-AvrLr41 preparations contained partially cleaved GST, yielding ∼85% and 70% purity for AvrLr41 and avrLr41_s692, respectively. Codon-optimized Lr41NLR^CC^ was fused with His-SUMO at its N terminus in the pET30a vector at the *Nde*I/*Hind*III sites and produced in DE3 cells in 1 L cultures with the same induction conditions as above. The protein was purified using a Ni column, followed by cleavage with the SUMO protease in the same elution buffer (50 mM Tris-HCl, 500 mM NaCl, pH 8.0). Recombinant purified Lr41NLR^CC^ with its SUMO tag removed was further purified on a Ni column and a Chromdex 75 column. For negative staining and cryo-EM analysis, individual proteins or protein mixtures were treated with TEV protease prior to sample preparation (1:1000 [v/v]; NEB#P8118, Australia).

### Surface plasmon resonance (SPR) analysis

SPR measurements were performed on a Biacore 8K instrument (Cytiva) using a CM5 sensor chip and HBS-EP+ running buffer (pH 7.4) (GenScript, Sydney, Australia) to quantify binding between recombinant purified Lr41 and GST-AvrLr41. Lr41 (with a His-tag; 0.18 mg/mL in 50 mM Tris-HCl, 150 mM NaCl, pH 8.0) was immobilized on flow cell 2 by EDC/NHS amine coupling after dilution with 10 mM sodium acetate (pH 5.0) and flow cell 1 served as the reference. Excess reactive esters were quenched with 1 M ethanolamine-HCl, and immobilization, activation, and quenching were performed following the manufacturer’s standard protocol. Recombinant purified GST-AvrLr41 (0.14 mg/mL in 50 mM Tris-HCl, 150 mM NaCl, pH 8.0) was serially diluted in running buffer (31.25-500 nM) and injected at 30 µL min^−1^ for 120 s of association and 180 s of dissociation, with 10 mM glycine-HCl (pH 1.5) used for surface regeneration between cycles. Sensorgrams were processed with Biacore 8K Evaluation Software v5.0 using reference-flow-cell subtraction and blank-buffer referencing; kinetic modeling employed the standard 1:1 Langmuir interaction framework without modification.

### Protoplast cell death assay

Protoplasts were isolated from 7-9-day-old wheat seedlings following Wilson et al. (2024)^43^. Briefly, leaves were scored and peeled to expose mesophyll cells, floated on top of 0.6 M mannitol (10 min), and digested in enzyme solution (20 mM MES pH 5.7, 0.8 M mannitol, 10 mM KCl, 10 mM CaCl_2_, 0.1% [w/v] BSA, 1.5% [w/v] cellulase R-10, 0.75% [w/v] macerozyme R-10) for 3 h with shaking at 60 rpm in darkness. Protoplasts were filtered, pelleted by centrifugation at 100 g for 3 min, washed in W5 buffer (154 mM NaCl, 125 mM CaCl_2_, 5 mM KCl, 2 mM MES, pH 5.7), counted, and resuspended to 3.5 × 10^5^/mL in MMG (4 mM MES, pH 5.7, 0.8 M mannitol, 15 mM MgCl_2_). Transfection reactions were performed in 96-well V-bottom plates using PEG-mediated uptake. Each reaction contained 50 µl protoplasts, 50 µl PEG solution (40% [w/v] PEG 4000, 0.2 M mannitol, 100 mM CaCl_2_), and plasmid DNA. For *Avr-R* co-expression, 0.5 µg of plasmid harboring *Avr*, 2 µg of plasmid carrying the *R* gene, and 5 µg of luciferase reporter plasmid were used; Avr-only controls used 1 µg of plasmid harboring *Avr* and 5 µg of the luciferase reporter plasmid. After 10 min of incubation, 200 µl W5 was added, and cells were incubated overnight at room temperature under dark conditions. For luciferase assays, protoplasts were lysed in 50 µl Promega Cell Lysis Buffer (#E3971) for 15 min, after which 50 µl lysate was transferred to opaque white 96-well plates (Corning #CLS3600). Baseline luminescence was measured (Tecan Infinite 200Pro, 1000 ms integration), followed by addition of 50 µl Steady-Glo substrate (Promega, #E2520). Luminescence was measured with and without green/red filters. Differences in normalized luminescence values were assessed using a one-way analysis of variance (ANOVA) followed by Tukey’s Honestly Significant Difference post hoc test.

The barley protoplast cell death assay was performed as previously described^61^ with modifications. Barley protoplasts were isolated from 10-day-old to 2-week-old ‘Golden Promise’ seedlings grown in greenhouse at 18°C. The abaxial epidermis of the leaves was removed with a razor to expose mesophyll cells; the abaxial side was placed in 0.6 M mannitol solution. The peeled leaves were transferred to the enzyme solution (10 mM MES, pH 5.7, 0.6 M mannitol, 20 mM KCl, 10 mM CaCl_2_, 0.9 M NaCl, 0.1% [w/v] BSA, 1.5% [w/v] cellulase R-10, 0.5% [w/v] macerozyme R-10) with shaking at 60 rpm in the dark for 3 h. Next, one volume of wash buffer (2 mM MES, pH 5.7, 5 mM KCl, 0.25 M CaCl_2_) was added to the cell suspension, and the protoplasts were filtered through 100-μm nylon mesh filters. The cells were collected by centrifugation at 100 g for 3 min. Wash buffer was added to the cells, and protoplasts were allowed to settle in the dark for 40 min. The supernatant was then removed and transfection buffer 1 (4 mM MES, pH 5.7, 0.4 M mannitol, 5 mM MgCl_2_) was added. The OD_600_ of pre-settled protoplasts was measured and adjusted to 0.4 if needed. For each transfection reaction, 8 μg of each plasmid was added to the protoplasts; in the case of multiple constructs, each construct was added at a 1:1:1 or 1:1 ratio (w/w/w or w/w). Transfection buffer 2 (0.2 M mannitol, 0.1 M CaCl_2_, 40% [w/v] PEG4000) was then added to the samples, the mixture inverted and incubated for 15 min before a second wash was performed. Before plating, because of slow accumulation of GFP signals, samples were resuspended in 500 μL regeneration buffer (4 mM MES, pH 5.7, 0.6 M mannitol, 20 mM KCl) and incubated overnight in the dark. Measurements were taken in a 96-well clear bottom black-walled plate (Falcon/Corning #353219) using a plate reader (Vantstar, BMG Labtech). The excitation/emission wavelengths for GFP were 470/15 nm and 515/20 nm, respectively. The plate was set to shake at 50 rpm for 10 s before each reading. The pGFPGUSplus plasmid and nontransfected protoplasts were used as internal and blank controls, respectively.

### BSMV-mediated gene expression and VIGS constructs and assays

All the sequences of the candidate genes were synthesized by the Integrated DNA Technologies (San Diego, CA, USA). For BSMV-mediated overexpression, full-length AvrLr41 without a putative small-secreted peptide was divided into three domains and each domain was cloned into the BSMV γ-vector. These candidate fragments were fused with a Ribosome Binding Site (RBS) and inserted in a reversed orientation after the stop codon of the BSMV gamma genome construct (γ-vector)^62^. *In vitro*-synthesized BSMV RNAs, including α, β, and γ-AvrLr41-1+ AvrLr41-2+ AvrLr41-3 (1:1:1 ratio) were pre-mixed and rub-inoculated onto the 2^nd^ leaf of the wheat *Lr41*+/- NILs at the 2-leaf stage. For *Lr39*/*Lr41* and *Lr41NLR^CC^* silence assays, each 150 bp (for *Lr39*/*Lr41*) and 205 bp (*Lr41NLR^CC^*) gene-specific fragment was inserted after the stop codon of the BSMV γ-vector. *In vitro*-synthesized BSMV RNAs, including α, β, and γ-target (1:1:1 ratio) as described in Huang (2017)^62^, were rub-inoculated onto the 2^nd^ leaf of wheat plants at the 2-leaf stage, with 20 seedlings per gene. Total RNAs from inoculated samples were extracted at 9 dpi for cDNA synthesis and qPCR. qPCR was carried out to check the expression of the genes relative to *TaActin* (table S9). For all virus experiments, at least five independent experiments were carried out. The presence of all fragments was detected by PCR in parallel to phenotypic assays, using the primers listed in table S9.

### Yeast two-hybrid (Y2H) assay

Protein-protein interactions were assessed using the Matchmaker Gold Y2H system (Takara Bio, Japan). Effector and NLR were cloned into the pGBKT7 and pGADT7 vectors, whereas the *Lr41* full-length coding sequence was cloned into pGADT7. The pGBKT7 vector and the resulting pGBKT7-based plasmids were introduced into the haploid strain Y187 and pGADT7 and its derived plasmids were introduced into the haploid strain Y2HGold. Variants of the *Lr41NLR* coding sequence were cloned into both vectors and the resulting plasmids introduced into both strains. Yeast strains harboring the plasmids of interest were mated overnight at 28 °C, with shaking at 200 rpm, and plated in ten-fold serial dilutions on SD medium lacking leucine and tryptophan, and on SD medium lacking leucine, adenine, histidine, and tryptophan and containing aureobasidin A and X-alpha-gal. Transformation and mating were performed according to the standard Matchmaker Gold protocol. Positive interactions were determined based on growth and reporter gene activation, indicated by blue colonies. Photographs were captured 3 days after plating.

### Agrobacterium infiltration in *N. benthamiana*

Agrobacterium (*Agrobacterium tumefaciens*) cultures were grown overnight at 28°C. Cultures with an OD of 1.0 were collected by centrifugation and resuspended to a final OD_600_ of 0.5 in MES buffer (10 mM MgCl_2_, 10 mM MES, pH 5.9) containing 300 μM acetosyringone. Greenhouse conditions were set to a 16-h light/8-h dark photoperiod at 25°C. The third and fourth leaves of 4-week-old *N. benthamiana* plants were infiltrated with 100 µL of Agrobacterium cell suspension into the abaxial leaf surface. Plants were incubated at 26 °C for two days in the dark and then under high humidity and low light conditions. Cell death was observed 3-5 days after infiltration.

### Protein extraction

Leaf discs were collected from the leaves of *N. benthamiana* plants at 1-3 days post-infiltration. Samples were flash-frozen in liquid nitrogen, ground with a mortar and pestle, and combined with 30 μL 4× Laemmli sample buffer (0.2 M Tris base, 0.28 M SDS, 40% [v/v] glycerol, 20% [v/v] 2-mercaptoethanol, 0.005% [w/v] bromophenol blue, pH 6.8), 24 μL DTT, and 66 μL dH_2_O, for immunoblot analysis. For co-immunoprecipitation (Co-IP), proteins were extracted using a gentle lysis buffer according to Duggan et al. (2021)^63^. Samples were vortexed for 90 s, denatured at 80 °C for 5 min, and centrifuged for 10 min at 16,000 rpm. The supernatant was run on a gel or stored at −20 °C for future use.

### Immunoblot, co-immunoprecipitation (Co-IP), and pull-down assays

Protein samples were resolved on 4-12% Bolt Bis-Tris Plus Mini Protein Gels (Thermo Fisher Scientific, NW04122BOX) at 200 V for 22 min in MES SDS buffer. Proteins were transferred to PVDF membranes in 1× transfer buffer substituted with methanol and Bolt antioxidant (Thermo Fisher Scientific, BT0005) at 20 V for 50 min. Prior to transfer, membranes were soaked in methanol, rinsed in water, and soaked in transfer buffer for 5 min. Prestained protein ladders (Thermo Fisher, #26620) were included for size estimation. All antibodies were reconstituted in 5% (w/v) non-fat dry milk with 1× Tris-buffered saline with 0.5% Tween-20 (TBST). Primary IgG antibodies (Thermo Fisher) were used at the following dilutions or concentrations: anti-HA 0.2 μg/mL (#26183), anti-GFP 1:3,000 (MA5-15256), anti-RFP 1:2,000 (MA5-15257), anti-GST 1:1,000 (MA4-004), anti-His 1:3,000 (MA1-21315). Membranes were blocked for 1 h at room temperature in 5% (w/v) non-fat dry milk in 1× TBST, and incubated with primary antibodies at 4°C overnight. Membranes were washed in 1× TBST three times for 10 min each time and incubated with secondary goat anti-mouse antibody (Thermo Fisher, #31430) at a dilution of 1:15,000 for 2 h at room temperature (with shaking). Membranes were then washed six times for 5 min each time and incubated with Pierce ECL substrate (Thermo Fisher, #32209) for 1 min. Images were captured with exposure times of 5-50 minutes using the BioRad ChemiDoc gel imaging system with chemiluminescent setting. For pull-down assays, a Thermo Classic Magnetic Co-IP kit (88804) was used. Protein extracts (100 μg) were incubated with 2 μg antibody, and bound complexes were captured on magnetic beads according to the standard kit protocol. All wash and incubation steps were performed according to the kit protocol, using the included buffers.

### Kinase assays

The kinase activity of Lr41 toward AvrLr41 was measured using a Universal Kinase Aassay Kit (Abcam, ab138879). Recombinant purified Lr41 was added at concentrations of 31.25, 62.5, 125, 250, and 500 nM; recombinant purified AvrLr41 and Lr41NLR^CC^ were kept at a constant concentration of 125 nM; ATP was kept constant at 100 μM. The standard curve was based on ADP concentrations of 0.05, 5, 10, 15, 20, and 30 μM. Before placing the plate in a plate reader, 20 μL of ADP sensor buffer and 10 μL of fluorescent ADP sensor were added to each well. Each sample was measured in triplicate. Measurements were taken in a clear bottom black-walled 96-well plate (Corning, Australia), every 5 min for 1.5 h using a plate reader (Vantstar, BMG Labtech). Measurements were taken in a spiral formation with excitation/emission wavelengths of 540/590 nm.

### Confocal microscopy

Leaf discs were harvested at 1-4 dpi, mounted in tap water under a coverslip, and imaged using 20× and 40× objectives under a Nikon ECLIPSE Ti2 inverted confocal microscope. ECFP settings were used for *AvrLr41* constructs, mRFP for *Lr41* constructs, and GFP as a control (pGFPGusPlus). Excitation wavelength for ECFP, mRFP, and GFP were 408 nm, 561 nm, and 490 nm, respectively, with emission wavelengths of 439-479 nm, 569-646 nm, and 488-552 nm, respectively. NIS elements software (v5.21.0) was used for visualization and editing images. Image processing included background subtraction, contrast adjustment, and advanced denoising based on automatic parameters.

### Negative staining

For negative staining, recombinant proteins were diluted 1:10 (v/v) and applied to positively glow-discharged 300-mesh copper grids. Glow discharge parameters were 25 mA for 30 s. Grids were washed twice with deionized water, briefly blotted dry and stained with uranyl acetate for 30 s. Finally, grids were briefly blotted dry and allowed to air-dry for at least 30 s. Micrographs were collected on an FEI Tecnai T12 (using a Gatan Rio 9 camera) operating with a tungsten filament with high tension set to 120 kV. Images were captured at 120-250× magnification.

### Cryo-electron microscopy (cryo-EM)

Purified protein samples were applied to Quantifoil Cu300 1.2/1.3 grids at a concentration of 200 µg/mL, with mixtures adjusted to a 1:1:1 (w/w/w) stoichiometric ratio where applicable. Grids for protein complexes were subjected to both a positive and negative glow discharge for 30 s at 25 mA; effector protein grids were negatively glow-discharged under the same conditions. Blotting was performed for 3 s at 100% humidity and 22 °C prior to plunge-freezing. Data were acquired under a Thermo Glacios microscope with a Ceta 16M CMOS camera over 16 h.

Image processing and 3D reconstruction were conducted using cryoSPARC (v4.7.1). Movies were processed in cryoSPARC using standard Patch Motion, Patch CTF, and particle-picking workflows.

EER movies were collected at 200 kV with a calibrated physical pixel size of 0.867 Å, a total exposure of 50 e^−^/Å^2^, spherical aberration of 2.7 mm, and an EER up-sampling factor of 2 with 40 fractions per movie. Motion correction was performed using Patch Motion (multi) with default settings, except for a maximum alignment resolution of 5 Å and a 500 Å^2^ alignment B-factor. CTF parameters were estimated using Patch CTF (multi) with default fitting settings, an amplitude contrast of 0.1, and a defocus search range of 1,000-40,000 Å. Initial particle identification was carried out using Blob Picker with a particle diameter range of 30-70 Å, circular blobs, and a 20 Å low-pass filter, followed by refinement of particle coordinates using Template Picker (templates low-pass filtered to 20 Å; angular sampling 5°). Up to 4,000 particle candidates per micrograph were retained. Particles were extracted with an extraction box size of 85 pixels using the default extraction workflow, applying a score threshold (min 0.157) and power-window filtering to exclude low-confidence picks. Extracted particles were then used for subsequent 2D/3D classification and refinement as described in Fig. S11f.

An initial atomic model of AvrLr41 was generated using the sharpened cryo-EM map in Phenix (v1.21). The model consisted of a single polypeptide chain (541 aa) with no ligand or water molecules. Manual model correction was performed using real-space refinement in Phenix, guided by backbone density and secondary-structure features of the sequence. Real-space refinement (RSR) was carried out, followed by model building and Dock in Map in Phenix (v1.21), using global minimization, atomic displacement parameter (ADP) refinement, Ramachandran restraints, rotamer restraints, and secondary-structure restraints. Refinement was iterated until convergence. The resulting model was further refined in Coot (v1.1.17) with overall RSR chain refinement with default restraints and secondary structure restraint parameters. Model geometry and stereochemistry were validated with MolProbity and Phenix validation tools. Map-model correlation coefficients were computed using masked and unmasked regions. Resolution estimates were taken from Phenix FSC-based procedures and reported statistics correspond to the final refined model and are listed in table S7.

## Materials availability

All the plasmids, rust, and plant materials generated and used in this study are available from the corresponding authors upon request under a material transfer agreement with The University of Sydney.

## Data and code availability

Raw sequencing reads were deposited in NCBI under project SUB15811007. The cryo-EM map of AvrLr41 was deposited in the EMDB database under accession number EMD-75312, and the atomic model was deposited in the PDB database under accession number 10NS. This paper does not report original code.

## Acknowledgements

We greatly acknowledge funding from the United States Department of Agriculture and the National Institute of Food and Agriculture (G179-24-WA310) to Y.D. and L.H.; the Judith and David Coffey and family to Y.D. and R.F.P.; the Grains Research and Development Corporation (GRDC: US221; US315; US00067; UOS1707-003RTX) to R.F.P. and P.Z., and the University of Sydney Strategic Research Fund (2024-G226587) to Y.D.. We thank Prof. Robert McIntosh for valuable and constructive suggestions. We thank Dr. Bhanu Mantri from the Transmission Electron Microscopy facility at the University of Sydney for technical assistance on cryo-EM sample preparation and TEM.

## Author contributions

C.L., E.N.A, E.P., B.D., S.S., T.N.H., R.F.P., P.Z. and Y.D. performed the experiments. E.N.A. and L.H. contributed to virus-related analyses. C.L., B.S., E.P., B.D., T.N.H., and E.L. contributed to plant protoplast analyses. D.X. and Y.D. performed the genomic and bioinformatic analyses. C.L., B.N. K., and Y.D. contributed to the microscopy analyses. C.L. performed the *N. benthamiana* assays. C.L. and Y.D. performed the protein biochemical analyses. A.K., J.M.A., and Y.D. performed the protein structural predictions. S.S. and P.Z. generated the EMS mutants. P. Z., L.H., and Y. D. performed the phenotypic analyses. R.F.P., P.Z., L.H., and Y.D. conceived and designed the project. Y.D. wrote the original draft. B.S., B.B.H.W., S.G.K., B.N.K., E.L., R.F.P., P.Z., L.H., and Y.D. reviewed and edited the manuscript. The authors declare no conflict of interest.

## Notes

### Competing Interest Statement

The authors have declared no competing interest.

## References

1. Ngou, B.P.M., Ding, P. & Jones, J.D.G. Thirty years of resistance: Zig-zag through the plant immune system. Plant Cell 34, 1447–1478 (2022).

2. Adachi, H., Derevnina, L. & Kamoun, S. NLR singletons, pairs, and networks: evolution, assembly, and regulation of the intracellular immunoreceptor circuitry of plants. Curr Opin Plant Biol 50, 121–131 (2019).

3. Wu, C.H., Derevnina, L. & Kamoun, S. Receptor networks underpin plant immunity. Science 360, 1300–1301 (2018).

4. Martin, R. et al. Structure of the activated ROQ1 resistosome directly recognizing the pathogen effector XopQ. Science 370(2020).

5. Yang, Y. et al. Paired plant immune CHS3-CSA1 receptor alleles form distinct hetero-oligomeric complexes. Science 383, eadk3468 (2024).

6. Forderer, A. et al. A wheat resistosome defines common principles of immune receptor channels. Nature 610, 532–539 (2022).

7. Bi, G. et al. The ZAR1 resistosome is a calcium-permeable channel triggering plant immune signaling. Cell 184, 3528–3541 e12 (2021).

8. Wang, J. et al. Ligand-triggered allosteric ADP release primes a plant NLR complex. Science 364(2019).

9. Wang, J. et al. Reconstitution and structure of a plant NLR resistosome conferring immunity. Science 364(2019).

10. Madhuprakash, J. et al. A disease resistance protein triggers oligomerization of its NLR helper into a hexameric resistosome to mediate innate immunity. Sci Adv 10, eadr2594 (2024).

11. Salmeron, J.M. et al. Tomato Prf is a member of the leucine-rich repeat class of plant disease resistance genes and lies embedded within the Pto kinase gene cluster. Cell 86, 123–33 (1996).

12. Shao, F. et al. Cleavage of Arabidopsis PBS1 by a bacterial type III effector. Science 301, 1230–3 (2003).

13. Wang, G. et al. The Decoy Substrate of a Pathogen Effector and a Pseudokinase Specify Pathogen-Induced Modified-Self Recognition and Immunity in Plants. Cell Host Microbe 18, 285–95 (2015).

14. Lewis, J.D. et al. The Arabidopsis ZED1 pseudokinase is required for ZAR1-mediated immunity induced by the Pseudomonas syringae type III effector HopZ1a. Proc Natl Acad Sci U S A 110, 18722–7 (2013).

15. Huard-Chauveau, C. et al. An atypical kinase under balancing selection confers broad-spectrum disease resistance in Arabidopsis. PLoS Genet 9, e1003766 (2013).

16. Reveguk, T. et al. Tandem kinase proteins across the plant kingdom. Nat Genet 57, 254–262 (2025).

17. Brueggeman, R. et al. The barley stem rust-resistance gene Rpg1 is a novel disease-resistance gene with homology to receptor kinases. Proc Natl Acad Sci U S A 99, 9328–33 (2002).

18. Klymiuk, V. et al. Cloning of the wheat Yr15 resistance gene sheds light on the plant tandem kinase-pseudokinase family. Nat Commun 9, 3735 (2018).

19. Chen, S. et al. Wheat gene Sr60 encodes a protein with two putative kinase domains that confers resistance to stem rust. New Phytol 225, 948–959 (2020).

20. Yu, G. et al. Aegilops sharonensis genome-assisted identification of stem rust resistance gene Sr62. Nat Commun 13, 1607 (2022).

21. Wang, Y. et al. An unusual tandem kinase fusion protein confers leaf rust resistance in wheat. Nat Genet 55, 914–920 (2023).

22. Cavalet-Giorsa, E. et al. Origin and evolution of the bread wheat D genome. Nature 633, 848–855 (2024).

23. Lu, P. et al. A rare gain of function mutation in a wheat tandem kinase confers resistance to powdery mildew. Nat Commun 11, 680 (2020).

24. Gaurav, K. et al. Population genomic analysis of Aegilops tauschii identifies targets for bread wheat improvement. Nat Biotechnol 40, 422–431 (2022).

25. Li, M. et al. A membrane associated tandem kinase from wild emmer wheat confers broad-spectrum resistance to powdery mildew. Nat Commun 15, 3124 (2024).

26. Arora, S. et al. A wheat kinase and immune receptor form host-specificity barriers against the blast fungus. Nat Plants 9, 385–392 (2023).

27. Jones, J.D.G., Staskawicz, B.J. & Dangl, J.L. The plant immune system: From discovery to deployment. Cell 187, 2095–2116 (2024).

28. Chen, R. et al. A wheat tandem kinase activates an NLR to trigger immunity. Science 387, 1402–1408 (2025).

29. Lu, P. et al. A wheat tandem kinase and NLR pair confers resistance to multiple fungal pathogens. Science 387, 1418–1424 (2025).

30. Upadhyaya, N.M. et al. Genomics accelerated isolation of a new stem rust avirulence gene-wheat resistance gene pair. Nat Plants 7, 1220–1228 (2021).

31. Chen, J. et al. Loss of AvrSr50 by somatic exchange in stem rust leads to virulence for Sr50 resistance in wheat. Science 358, 1607–1610 (2017).

32. Salcedo, A. et al. Variation in the AvrSr35 gene determines Sr35 resistance against wheat stem rust race Ug99. Science 358, 1604–1606 (2017).

33. Arndell, T. et al. Pooled effector library screening in protoplasts rapidly identifies novel Avr genes. Nat Plants 10, 572–580 (2024).

34. Deng, C. et al. The RppC-AvrRppC NLR-effector interaction mediates the resistance to southern corn rust in maize. Mol Plant 15, 904–912 (2022).

35. Chen, G. et al. Cloning southern corn rust resistant gene RppK and its cognate gene AvrRppK from Puccinia polysora. Nat Commun 13, 4392 (2022).

36. Shen, S. et al. Puccinia triticina avirulence protein AvrLr21 directly interacts with wheat resistance protein Lr21 to activate wheat immune response. Commun Biol 7, 1170 (2024).

37. Savary, S. et al. The global burden of pathogens and pests on major food crops. Nat Ecol Evol 3, 430–439 (2019).

38. Kolmer, J.A. GENETICS OF RESISTANCE TO WHEAT LEAF RUST. Annu. Rev. Phytopathol. 34(1996).

39. Huerta-Espino, J. et al. Global status of wheat leaf rust caused by Puccinia triticina. Euphytica 179, 143–160 (2011).

40. Cox, T.S., Raupp, W.J. & Gill, B.S. Leaf RustLResistance Genes Lr41, Lr42, and Lr43 Transferred from Triticum tauschii to Common Wheat. Crop Science 34, 339-343 (1994).

41. Singh, S. et al. Lr41, Lr39, and a leaf rust resistance gene from Aegilops cylindrica may be allelic and are located on wheat chromosome 2DS. Theor Appl Genet 108, 586–91 (2004).

42. Sun, X., Bai, G. & Carver, B.F. Molecular markers for wheat leaf rust resistance gene Lr41. Molecular Breeding 23, 311–321 (2008).

43. Wilson, S. et al. Multiplexed effector screening for recognition by endogenous resistance genes using positive defense reporters in wheat protoplasts. New Phytol 241, 2621–2636 (2024).

44. Kwon, A. et al. Tracing the origin and evolution of pseudokinases across the tree of life. Sci Signal 12(2019).

45. Seong, K. & Krasileva, K.V. Prediction of effector protein structures from fungal phytopathogens enables evolutionary analyses. Nat Microbiol 8, 174–187 (2023).

46. Hewitt, T. et al. Wheat leaf rust resistance gene Lr13 is a specific Ne2 allele for hybrid necrosis. Mol Plant 14, 1025–1028 (2021).

47. Sung, Y.C. et al. Wheat tandem kinase RWT4 directly binds a fungal effector to activate defense. Nat Genet 57, 1238–1249 (2025).

48. Baudin, M., Schreiber, K.J., Martin, E.C., Petrescu, A.J. & Lewis, J.D. Structure-function analysis of ZAR1 immune receptor reveals key molecular interactions for activity. Plant J 101, 352–370 (2020).

49. Bai, S. et al. Structure-function analysis of barley NLR immune receptor MLA10 reveals its cell compartment specific activity in cell death and disease resistance. PLoS Pathog 8, e1002752 (2012).

50. Cesari, S. et al. Cytosolic activation of cell death and stem rust resistance by cereal MLA-family CC-NLR proteins. Proc Natl Acad Sci U S A 113, 10204–9 (2016).

51. Raupp, W.J., Sukhwinder, S., Brown-Guedira, G.L. & Gill, B.S. Cytogenetic and molecular mapping of the leaf rust resistance gene Lr39 in wheat. Theoretical and Applied Genetics 102, 347–352 (2001).

52. Wu, C.H. et al. NLR network mediates immunity to diverse plant pathogens. Proc Natl Acad Sci U S A 114, 8113–8118 (2017).

53. Huang, N. & Li, H. compleasm: a faster and more accurate reimplementation of BUSCO. Bioinformatics 39(2023).

54. Love, M.I., Huber, W. & Anders, S. Moderated estimation of fold change and dispersion for RNA-seq data with DESeq2. Genome Biol 15, 550 (2014).

55. Adachi, H. et al. An N-terminal motif in NLR immune receptors is functionally conserved across distantly related plant species. Elife 8(2019).

56. Trapnell, C. et al. Differential analysis of gene regulation at transcript resolution with RNA-seq. Nat Biotechnol 31, 46–53 (2013).

57. Wu, J.Q. et al. A Chromosome-Scale Assembly of the Wheat Leaf Rust Pathogen Puccinia triticina Provides Insights Into Structural Variations and Genetic Relationships With Haplotype Resolution. Front Microbiol 12, 704253 (2021).

58. Vogt, F., Shirsekar, G. & Weigel, D. vcf2gwas: Python API for comprehensive GWAS analysis using GEMMA. Bioinformatics 38, 839–840 (2022).

59. Schwessinger, B. et al. A Near-Complete Haplotype-Phased Genome of the Dikaryotic Wheat Stripe Rust Fungus Puccinia striiformis f. sp. tritici Reveals High Interhaplotype Diversity. mBio 9(2018).

60. Van Dongen, S. Graph Clustering Via a Discrete Uncoupling Process. SIAM Journal on Matrix Analysis and Applications 30, 121–141 (2008).

61. Saur, I.M.L., Bauer, S., Lu, X. & Schulze-Lefert, P. A cell death assay in barley and wheat protoplasts for identification and validation of matching pathogen AVR effector and plant NLR immune receptors. Plant Methods 15, 118 (2019).

62. Huang, L. BSMV-Induced Gene Silencing Assay for Functional Analysis of Wheat Rust Resistance. Methods Mol Biol 1659, 257–264 (2017).

63. Duggan, C. et al. Dynamic localization of a helper NLR at the plant-pathogen interface underpins pathogen recognition. Proc Natl Acad Sci U S A 118(2021).

