## Supplemental figures for "Direct effector recognition by a tandem kinase triggers non-canonical immunity in wheat"

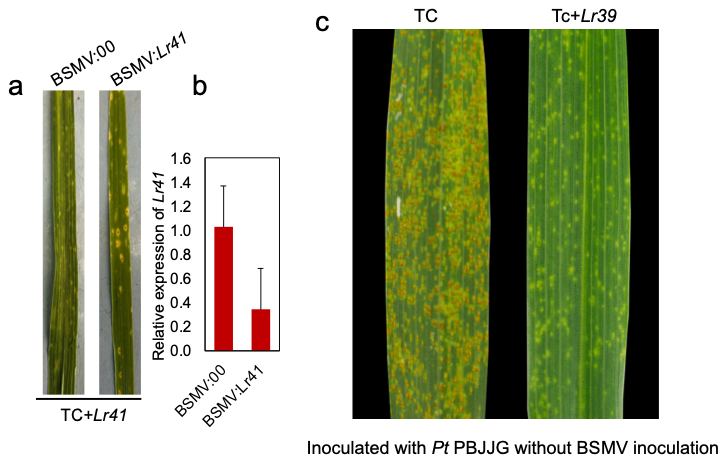

Fig. S1 (a) Virus-induced gene silencing of *Lr41*. Plants were challenged with *P. triticina* pathotype PBJJG. (b) Relative *Lr41* expression levels were measured by RT-qPCR. A non-silencing BSMV:00 was used as a control. Representative phenotype of two independent experiments were shown. (c) Thatcher (Tc) and the Tc+*Lr39* near-isogenic line inoculated with *P. triticina* pathotype PBJJG avirulent on *Lr41*.

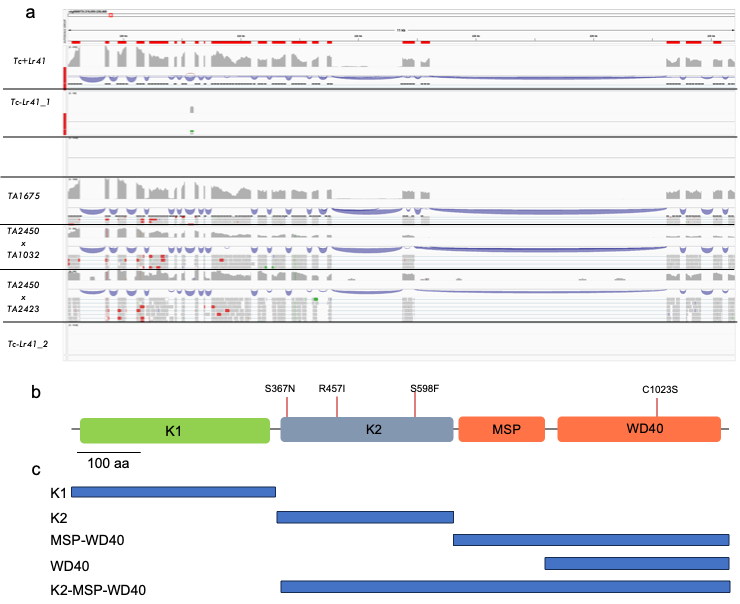

Fig. S2 (a) RNA-seq reads mapping to the TA1675 reference genome of the *Lr41* locus comparing with Tc+*Lr41*, three replicates of Tc, and four different *Ae. tauchii* lines. (b) Schematic overview of Lr41 protein domains with EMS-derived mutation locations (S367N and S598F) and haplotype-variants (R457I and C1023S) labelled. (c) Schematic overview of Lr41 protein regions tested in the Y2H analyses.

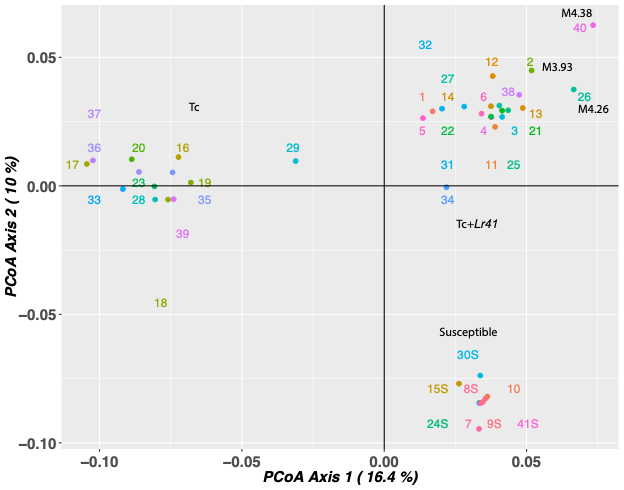

Fig. S3 Screening of *Lr41* EMS-derived mutant by Diversity Array Technology sequencing (DArTseq). Seeds of each individual were directly subjected to gDNA extraction for DArTseq. Principal component analysis (PCoA) differentiated M3 and M4 generation of the Tc+*Lr41* mutants from susceptible individuals and Tc background (Suppl. table 2). Three mutants (M3.93, M4.26 and M4.38) were selected for additional sequencing confirmation.

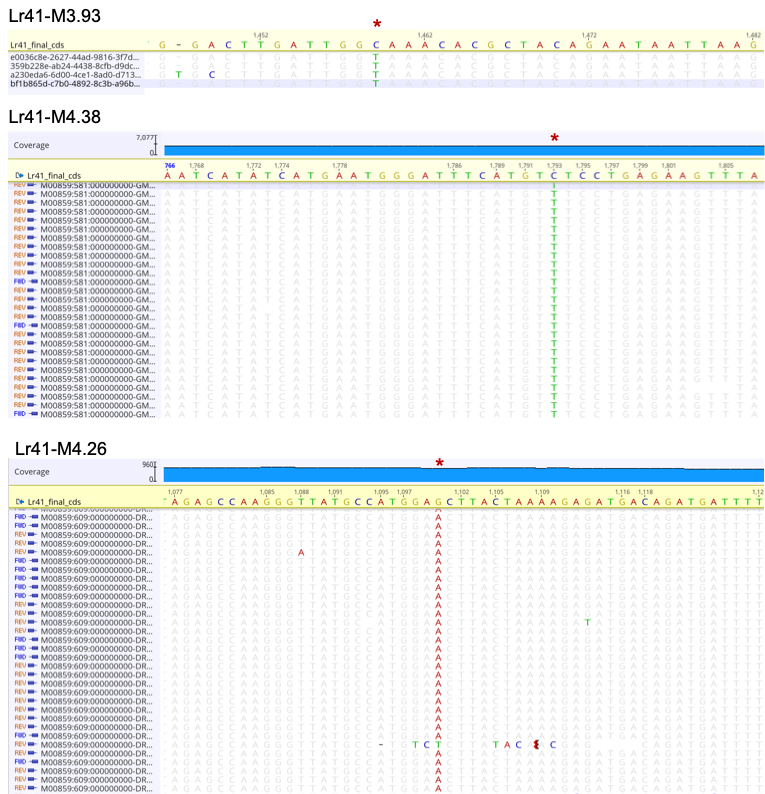

Fig. S4 Sequencing confirmation by Oxford Nanopore for the M3 generation mutant Lr41-M3.93 and Illumina for the M4 mutants Lr41-M4.38 and M4.26. Reads were mapped to the *Lr41* CDS sequence as a reference, and SNPs in the EMS mutants were highlighted.

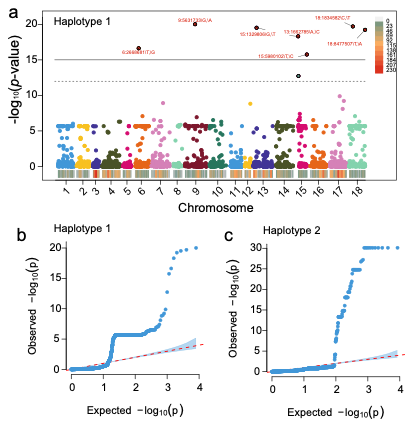

Fig. S5 Haplotype-dependent GWAS for *Lr41* avirulence. (a) Manhattan plot from the haplotype 1-based association analysis (using the s473 haplotype A as reference), showing dispersed SNP signals of modest significance across the chromosomes. Two horizontal lines mark the genome-wide significance thresholds at *p* = 1×10⁻¹² and *p* = 1×10^-15^. (b) Quantile-quantile (QQ) plot for the haplotype-1 GWAS analysis displaying strong upward deviation from the null distribution not attributable to a single association peak. (c) QQ plot for the haplotype 2-based GWAS analysis. Pronounced upper-tail enrichment was observed likely corresponding to chromosome 15 associated peak at the *AvrLr41* locus.

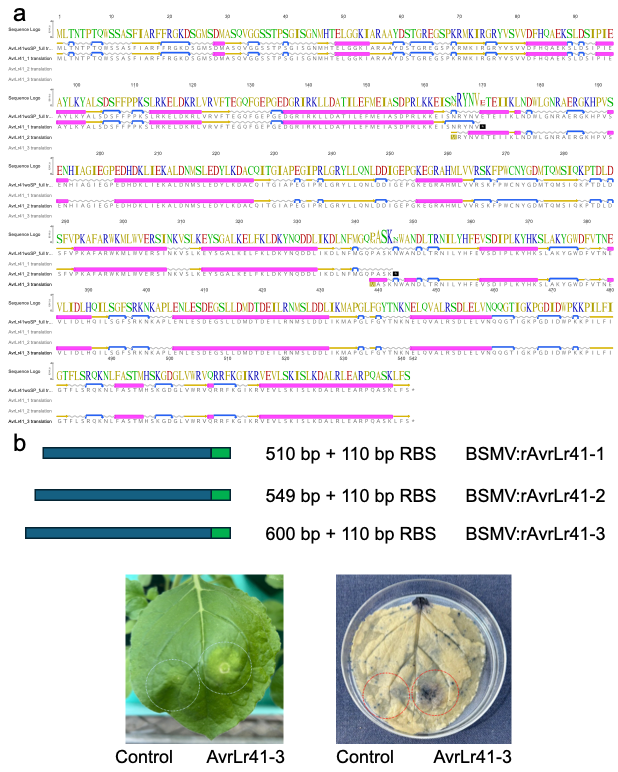

Fig. S6 Designing of fragments of *AvrLr41* for BSMV-mediated overexpression. (a) AA sequence alignment of full-length AvrLr41 along with the three fragments, ArvLr41-1, -2 and -3. AvrLr41-2 partially overlapped with both AvrLr41-1 and -3. Breaks were selected based on the secondary structure prediction in coil linking regions. Pink: alpha-helix; yellow: beta-sheet; grey: coil; blue: turn. (b) Schematic representation of all three fragments of *AvrLr41* used to assemble BSMV:*rAvrLr41-1*, BSMV:*rAvrLr41-2* and BSMV:*rAvrLr41-3*.

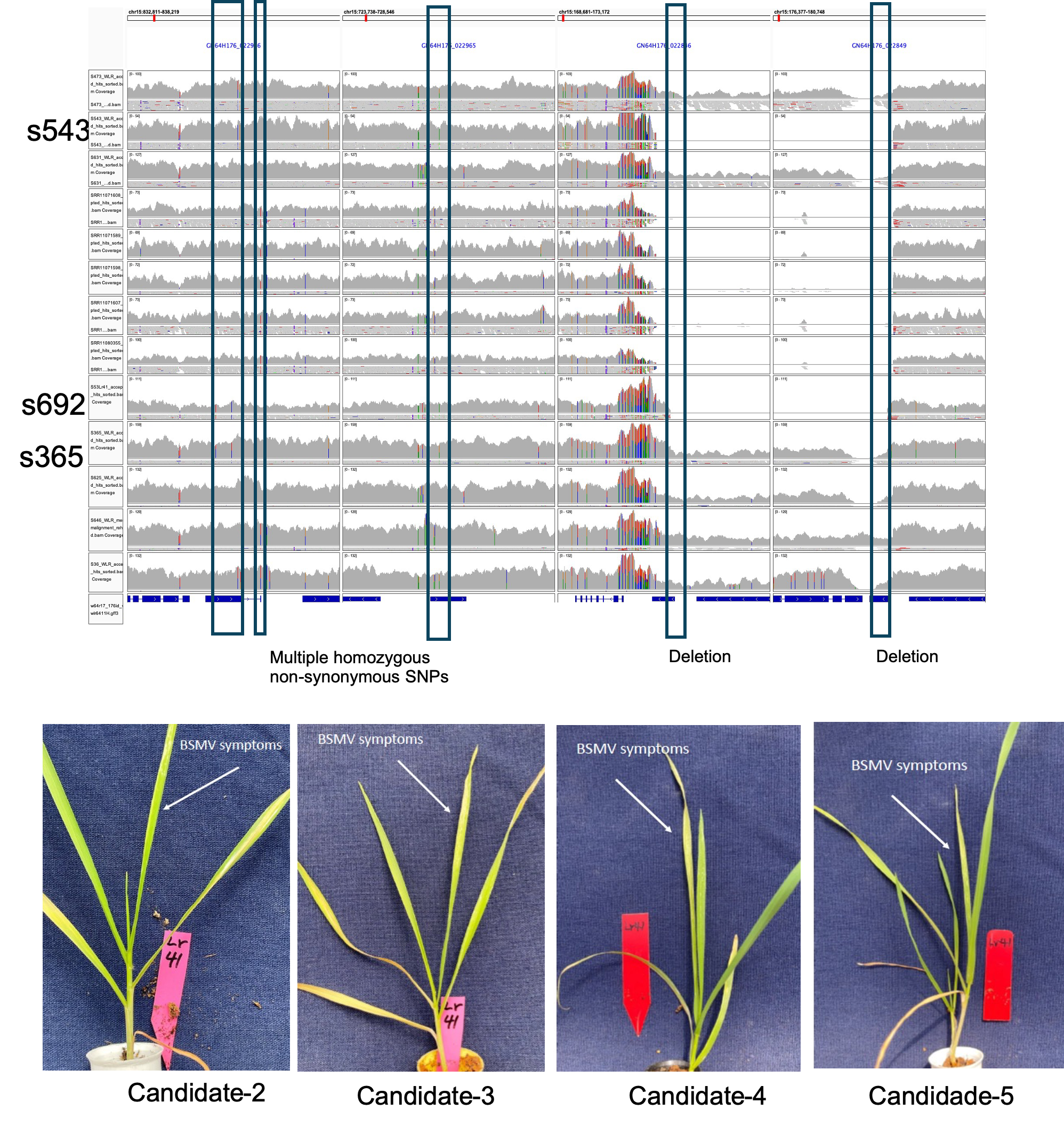

Fig. S7 Validation of AvrLr41 candidates. (a) Overview of additionally identified AvrLr41 candidates annotated in the loss of heterozygosity and GWAS top peak region. Variants were found within this region in the single-step greenhouse mutant s692 (*Lr41*-virulent) derived from s365 (*Lr41*-avirulent), as well as all international isolates (5 were shown) with reads coverage over 20x. (b) BSMV-mediated overexpression of the AvrLr41 candidates in Tc+*Lr41* showed no change in BSMV viral symptoms.

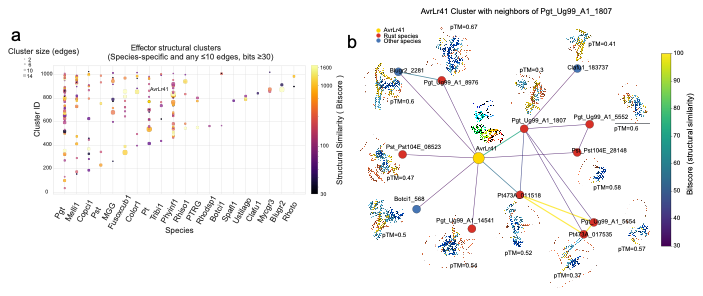

Fig. S8 Structural organization and molecular networking of effector families reveals a discrete AvrLr41-centered cluster with marginal structural resemblance to other plant fungal or oomycete pathogen small secreted proteins. (a) Species-specific effector structural clusters with less than 10 edges and FoldSeek bitscore over 30. Scatter plot showing structural similarity clusters recovered from all-against-all FoldSeek comparisons across 20 fungal and oomycete species. A total of 30,218 predicted protein structures were compared, generating 3,849,135 pairwise structural alignments and an initial 1,785 structural clusters. Clusters that are species-unique and have ≤10 total similarity edges were retained. A refined set of 207 clusters remained for visualization. Among these, 76 clusters belong to rust pathogens (*Puccinia graminis, P. triticina,* and *P. striiformis*; 16 correspond to *P. triticina* including AvrLr41), and 131 are from other fungal or oomycete species. X-axis denotes species inferred from protein prefixes and Y-axis denotes cluster ID. Bubble size represents connectivity within each cluster (number of structural similarity edges). Color scale represents structural similarity as bitscore of all pairwise comparisons within the cluster. AvrLr41 is highlighted within the *P. triticina* lineage. (b) AvrLr41-centered structural similarity network and its nearest neighbors. Network visualization of all proteins with direct structural similarity to AvrLr41 (FoldSeek bitscore ≥30). Nodes represent effector proteins; edges represent pairwise structural alignments; edge colors represent alignment bitscore. Node color indicates species group: yellow, AvrLr41; red, rust pathogens (*P. graminis, P. triticina,* and *P. striiformis*); blue, non-rust species. AF3 structures are shown for each neighbor with predicted TM-score (pTM). Edges connecting AvrLr41 to Pgt_Ug99_A1_1807 followed by Pt473A_011518 showed the highest bitscores. Cryo-EM resolved structure model of AvrLr41 was used.

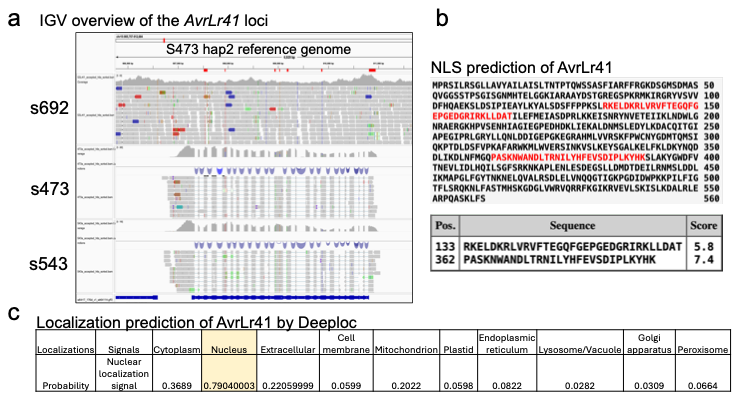

Fig. S9 Sequence feature overview and nuclear-localization predictions for AvrLr41. (a) Integrative Genomics Viewer (IGV) overview of the *AvrLr41* locus aligned to the s473 haplotype-2 reference genome. Short-read whole-genome sequencing and mRNA-seq alignments for isolates s692, s473, and s543 were shown. Exon-intron boundaries of *AvrLr41* are indicated by the mRNA-seq reads mapping coverage and reference annotation, with splicing junctions labelled. Read coverage tracks and SNPs highlight the sequence differences present in the virulent isolates (s692 and s543, homozygous SNPs) and avirulent/reference isolate s473 (showing exclusively heterozygous SNPs) relative to the reference (s473) genome. (b-c) Nucleus localisation prediction could be consistently obtained for AvrLr41. (b) Classical nuclear localization signal (NLS) predictions for AvrLr41 using cNLS Mapper. Two NLS motifs were predicted and highlighted in red within the primary sequence. The corresponding cNLS Mapper scores (5.8 and 7.4) were shown under a positive thread of 5.0, along with the position of each predicted motif. (c) DeepLoc-1.0 subcellular localization predictions for AvrLr41. Probability outputs for all modelled compartments are tabulated; the highest-probability assignment corresponds to nuclear localization.

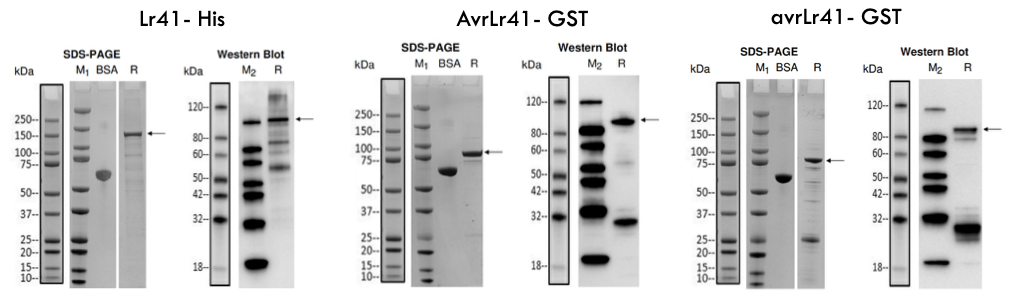

Fig. S10 Recombinant expression and quality control of Lr41-His, AvrLr41-GST, and avrLr41-GST proteins. Lr41-His was heterologously expressed in *E. coli* BL21 Star™ (DE3) inducted by 0.5 mM IPTG at 15 °C for 16 h, and analysed by SDS–PAGE and Western blot with anti-His antibody. Codon-optimized AvrLr41-GST and avrLr41-GST were expressed in *E. coli* Rosetta2 (DE3) at 16 °C for 16 h following 0.5 mM IPTG induction. For each construct, soluble fractions (R) were detected by SDS-PAGE alongside molecular weight markers (M₁; Bio-Rad, 1610374S) and BSA controls. For Western blot a different ladder was used (M_2_; GenScript, M00673) and probed with anti-GST or anti-His antibodies. Arrows indicate the recombinant proteins at their expected molecular masses.

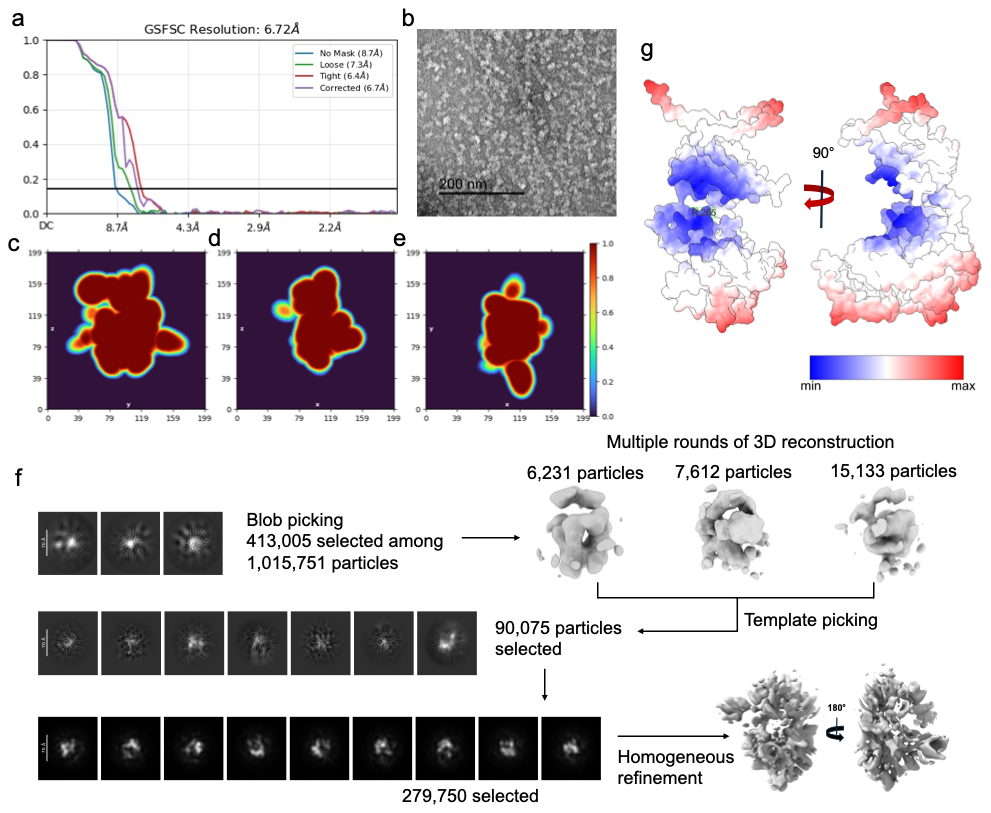

Fig. S11 Cryo-EM reconstruction and quality assessment of AvrLr41. (a) Gold-standard Fourier shell correlation (GS-FSC) curves for the single-particle cryo-EM reconstruction of AvrLr41 using CryoSPARC (v4.7.1), calculated without a mask and with loose, tight, and mask-corrected masks as indicated. The mask-corrected FSC curve crosses the 0.143 criterion at 6.7 Å, which is taken as the overall map resolution (unmasked, loose, and tight masks give 8.7, 7.3, and 6.4 Å, respectively). (b) Representative electron micrograph of negatively stained AvrLr41 particles showing a homogeneous monodisperse particle distribution (scale bar, 200 nm). (c–e) Directional FSC analysis of the final map, shown as 2D projections of the 3D FSC volume along the z, x, and y axes, respectively. Color scale (0-1) reports FSC values, with warm colors corresponding to higher correlation. (f) Workflow of particle selection and 3D reconstruction in CryoSPARC. Since particles were small, they were initially obtained from blob picking and were first 2D-classified and subsequently used to generate an initial 3D model. This initial volume was then used as a template for template-based particle picking, followed by additional rounds of 2D cleaning and 3D reconstruction. Iterative refinement of progressively improved particle sets produced the final homogeneous reconstruction shown at right (180° views). (g) Surface representation of the refined AvrLr41 model colored by atomic B-factors, showing relative mobility and map support across the structure. Lower B-factors (blue) correspond to the regions of stronger local density and higher confidence in model placement, whereas higher B-factors (red) indicate the areas of reduced density, increased flexibility, or limited structural information at the achieved resolution. Two orthogonal views (90° rotation) highlight that the central core of the protein is well supported by the cryo-EM map, while peripheral domains exhibit increased disorder consistent with map attenuation at ~6 Å resolution. A continuous B-factor gradient across the surface demonstrates that the refined model fits the available density without over-sharpening or localized overfitting.

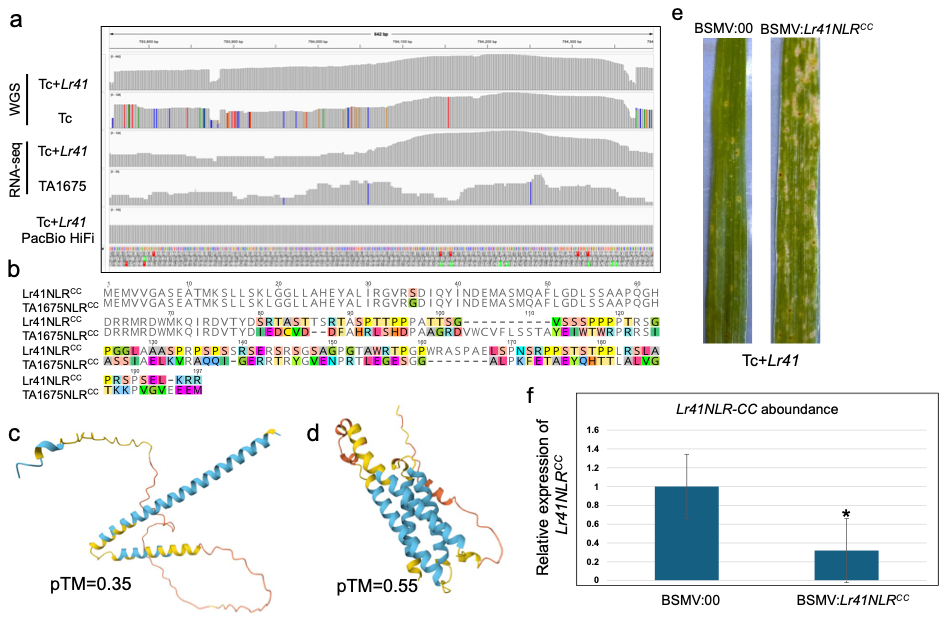

Fig. S12 Comparative genomic, sequence, and structural analyses of the Lr41-associated truncated NLR (Lr41NLR^CC^) and its intact *Ae. tauschii* ortholog (AetNLR). (a) IGV visualization of short-read genomic, RNA-seq and PacBio HiFi long-read alignments across the *Lr41NLR^CC^* locus in Tc+*Lr41* and the corresponding intact *NLR* locus in *Ae. tauschii* TA1675 as compared to Tc. Tc+*Lr41* genome assembly was used as a reference. (b) Pairwise aa alignment of Lr41NLR^CC^ and AetNLR^CC^ highlighting conserved N-terminal CC motifs and the loss of the third and fourth CC helices in Lr41NLR^CC^. Polymorphic residues (colour-coded) reflect divergence at both conserved and low-complexity positions. (c–d) AlphaFold3 structural predictions of Lr41NLR^CC^ (c) and AetNLR^CC^ (d). Lr41NLR^CC^ displayed as a two-helix CC bundle followed by an extended intrinsically disordered C-terminal region. By contrast, AetNLR adopts the canonical four-helix CC fold expected for full-length CNL-type NLRs. (e) Phenotypic assay of Tc+*Lr41* plants following BSMV-induced silencing of *Lr41NLR^CC^*. Tc+*Lr41* plants inoculated with the BSMV:00 control exhibited the characteristic *Lr41*-specific resistance to the *P. triticina* race PBJJG, with only very few, small pustules consistent with a typical HR response. By contrast, VIGS of *Lr41NLR^CC^* (BSMV:*Lr41NLR^CC^*) resulted in reduction in resistance, with clearly visible uredinial pustules indicating partial compromise of *Lr41*-mediated defense. (f) Relative expression of *Lr41NLR^CC^* in Tc+*Lr41* plants following VIGS treatment. qRT-PCR analysis confirmed high transcript abundance in BSMV:00 control plants, whereas *Lr41NLR^CC^* expression was substantially reduced in BSMV:*Lr41NLR^CC^* plants, consistent with effective silencing and the observed attenuation of resistance (Student’s t-test, **p*<0.05). Shown as representative of at least 20 plants and 3 independent experiments.

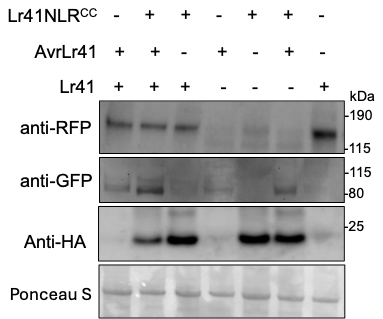

Fig. S13 Protein expression and Western blot for Lr41NLR^CC^, AvrLr41 and Lr41. Samples collected in parallel for HR confocal and HR assays (as displayed in Fig. 4c) were harvested at 1 dpi.

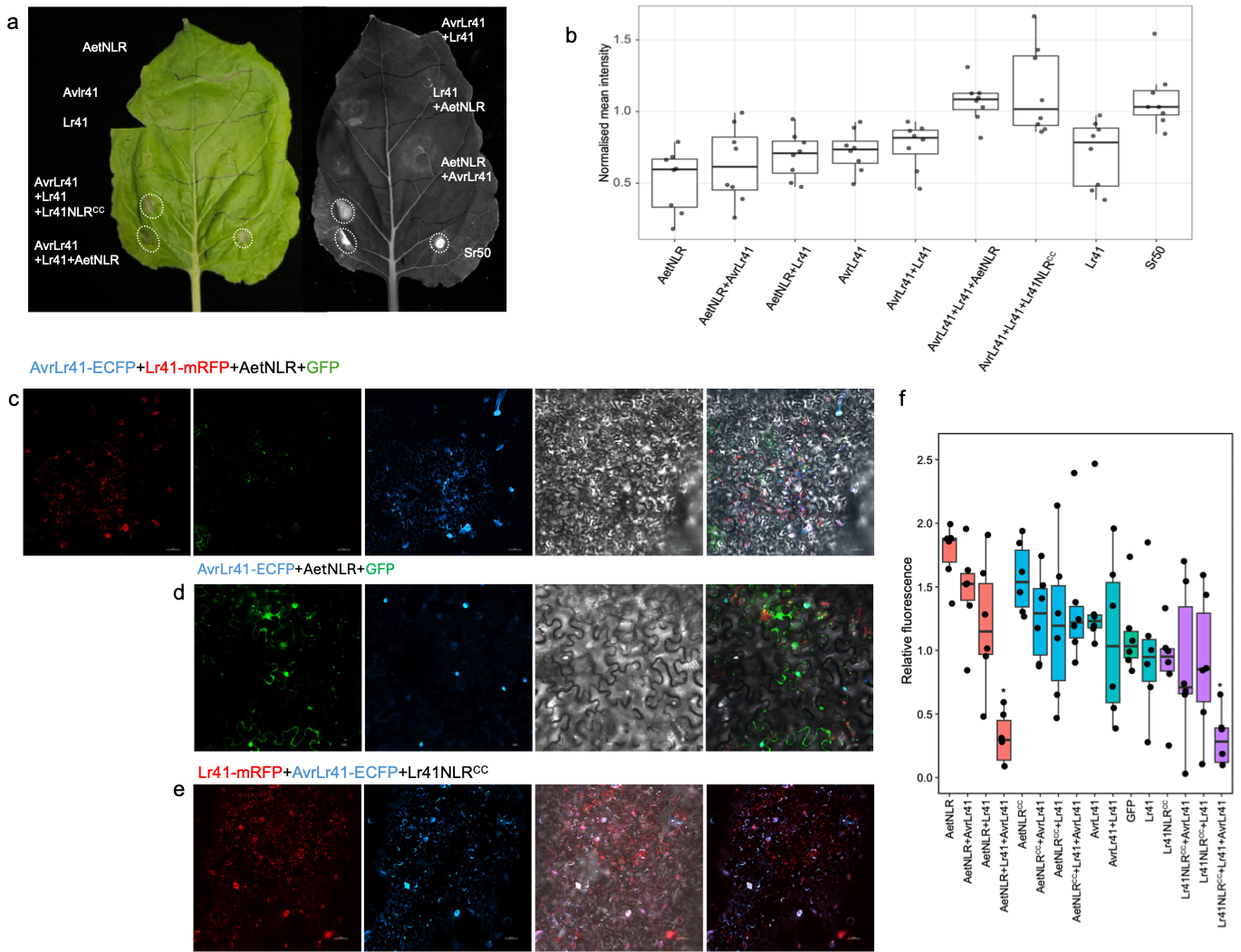

Fig. S14 AetNLR can substitute for the truncated Lr41NLR^CC^ in Lr41-AvrLr41-dependent immunity in *Nicotiana benthaminana* and barley protoplasts. (a) *N. benthamiana* co-expression assays assessing helper NLR requirements for *Lr41*-dependent HR responses. Agrobacterium expressing AvrLr41, Lr41, and either the truncated Lr41NLR^CC^ or the full-length AetNLR were infiltrated in the combinations indicated. HR occurred only when a helper NLR (Lr41NLR^CC^ or AetNLR) and both recognition components (Lr41 and AvrLr41) were co-expressed. Neither individual constructs nor incomplete combinations elicited HR. Sr50 served as a positive control. Circles denote representative HR from ≥6 independent infiltrations per condition. A representative photo taken at 3 dpi was shown. (b) Quantification of HR-associated autofluorescence (normalized mean intensity) from independent infiltration sites (n≥6 per combination). Data are shown as individual points. (c) Confocal microscopy of epidermal cells co-expressing AvrLr41-ECFP, Lr41-mRFP, AetNLR-HA at 1 dpi. 35S:GFP expression was combined to check the expression site. Co-expression resulted in punctate and peripheral accumulation of all three proteins, consistent with a multi-component complex. Channels: mRFP (red), GFP (green), ECFP (blue), bright-field, and merged composite. Scale bars, 10 μm. (d) Confocal images of cells expressing AetNLR, AvrLr41-ECFP and free GFP without Lr41. No HR or characteristic peripheral co-localization was observed, indicating that AetNLR alone is insufficient to relocalize AvrLr41 or trigger defense. (e) Confocal microscopy images of epidermal cells co-expressing AvrLr41-ECFP, Lr41-mRFP, Lr41NLR^CC^ at 2 dpi. *35S:GFP* expression was combined to check the expression site. Co-expression resulted in punctate and peripheral accumulation of all three proteins, consistent with a multi-component complex. Scale bars, 10 μm. (f) Change in relative fluorescence in barley (cv. Golden Promise) protoplasts following transient (co)expression of the indicated constructs. Protoplasts were transfected and pre-incubated overnight to allow recovery. Relative fluorescence values represent background-corrected fluorescence at 6 h and 16 h, respectively. Data are shown as boxplots with individual data points overlaid; boxes indicate the interquartile range, centre lines indicate the median. Statistical significance was assessed using a Kruskal-Wallis test followed by Dunn’s multiple comparisons test with Holm adjustment. Asterisks indicate significant differences relative to GFP control (**p* < 0.05). Two independent biological replicates were performed each with three technical replicates.

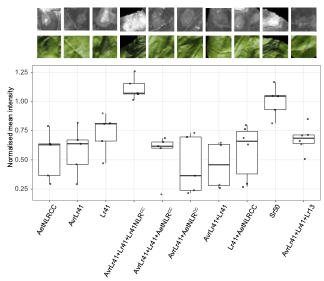

Fig. S15 Neither AetNLR^CC^ nor Lr13 trigger HR together with Lr41 and ArvLr41.

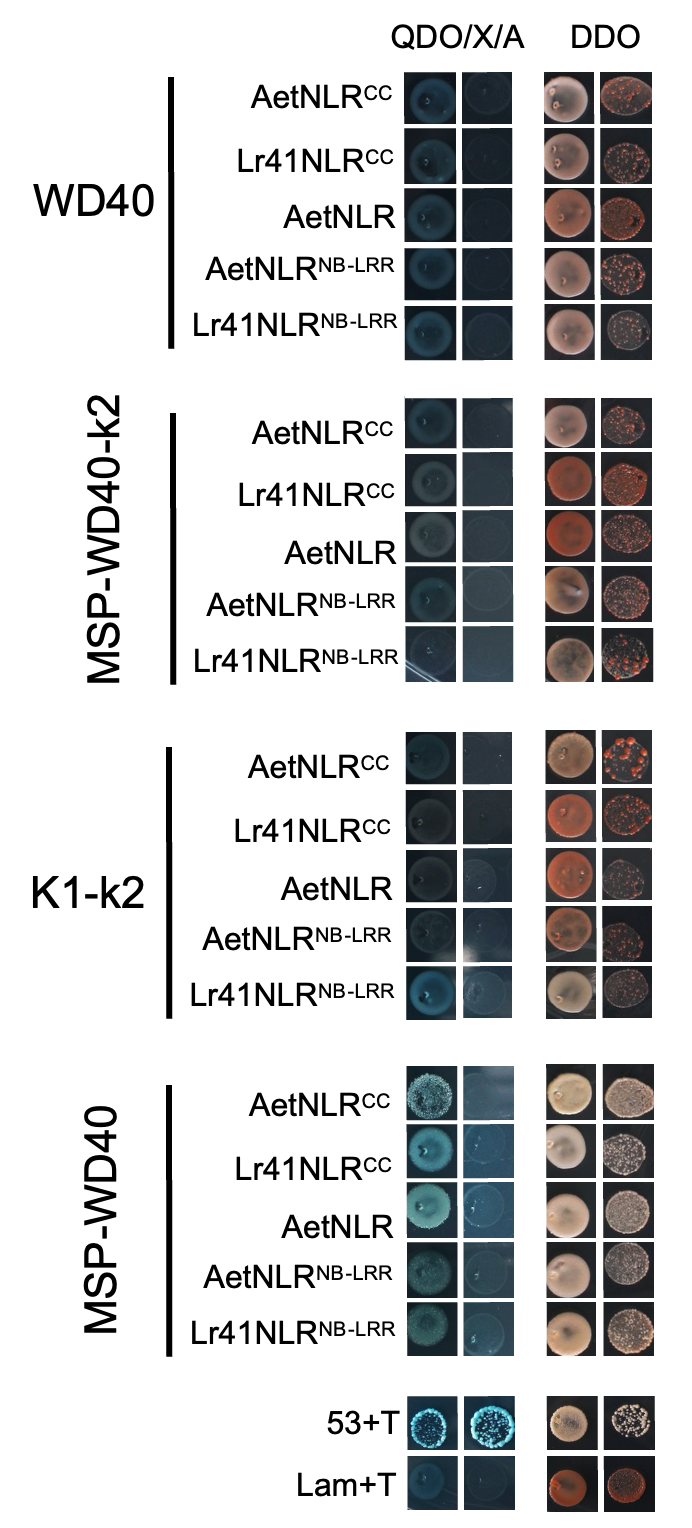

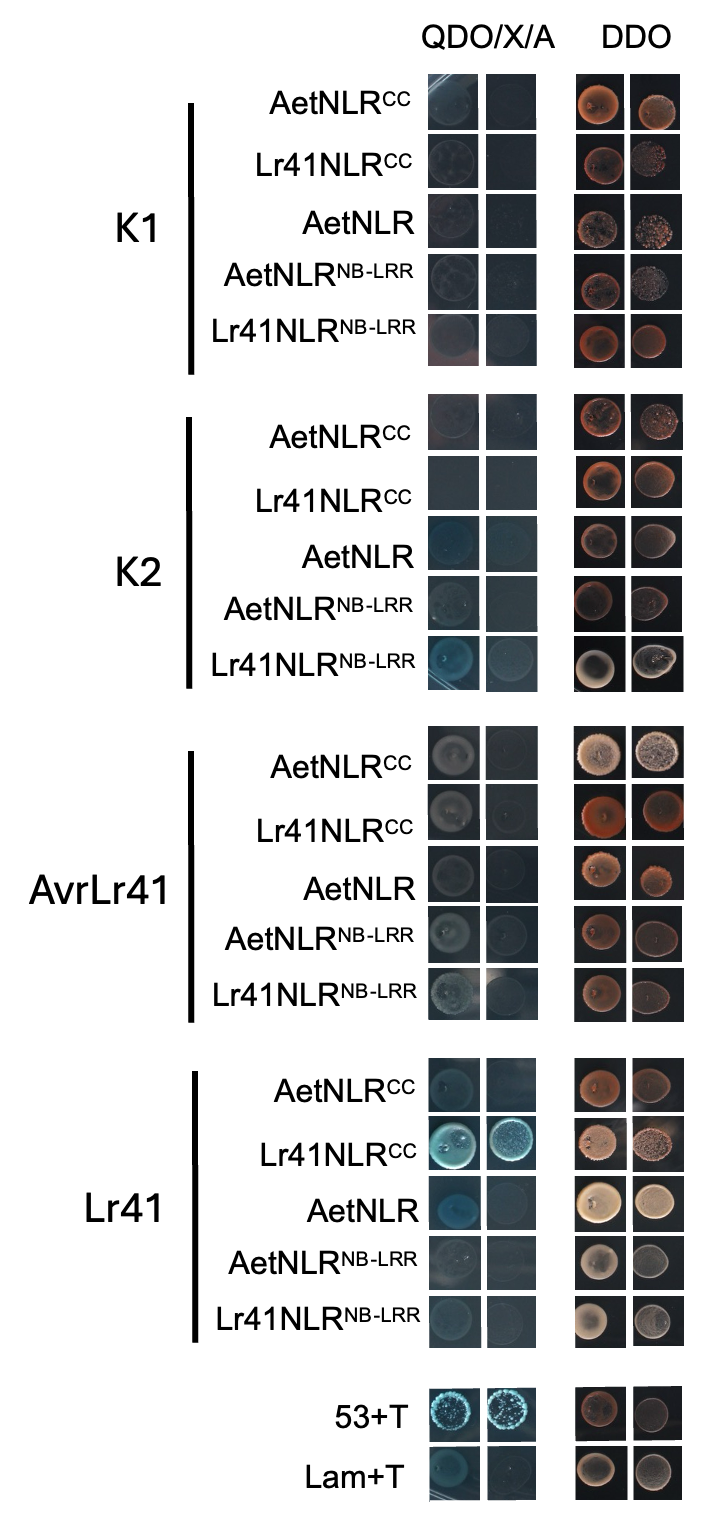

Fig. S16 Lr41 directly interacts with Lr41NLR^CC^ but not with AetNLR and its variant. MSP-WD40 domain showed weak interaction with all helper NLR versions. Constructs fused to the GAL4 activation domain (AD) were paired with constructs fused to the GAL4 DNA-binding domain (BD), transformed into yeast, and selected on SD/–Leu/–Trp (DDO) medium prior to interaction testing on stringent SD/–Leu/–Trp/–His/–Ade medium supplemented with X-α-gal and Aureobasidin A (QDO/X/A). Growth and blue coloration indicate positive interaction. Representative results of three independent experiments were shown.

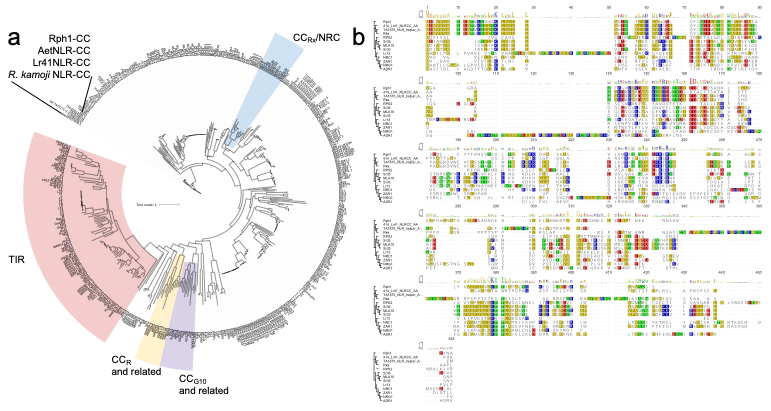

Fig. S17 Phylogeny and sequence analyses of Lr41-associated helper NLRs. (a) Maximum-likelihood phylogeny of 420 plant NLR N-terminal regions, each truncated to the first 250 aa, including CC-type NLRs and TIR-type NLRs, along with Lr41NLR^CC^ and its homologs from *Roegneria kamoji* and *Ae. tauschii*. The tree resolves the major CC-domain lineages, including the TIR-type outgroup (pink), CC_R_ and related CC subclasses (yellow), CC_G10_ and related CC subclasses (purple), and the CC_RX_/NRC superclade (blue). Lr41NLR^CC^ and AetNLR clustered distinctly in separate branches and close to the barley Rph1^CC^. (b) Multiple sequence alignment of representative CC domains from each highlighted clade.

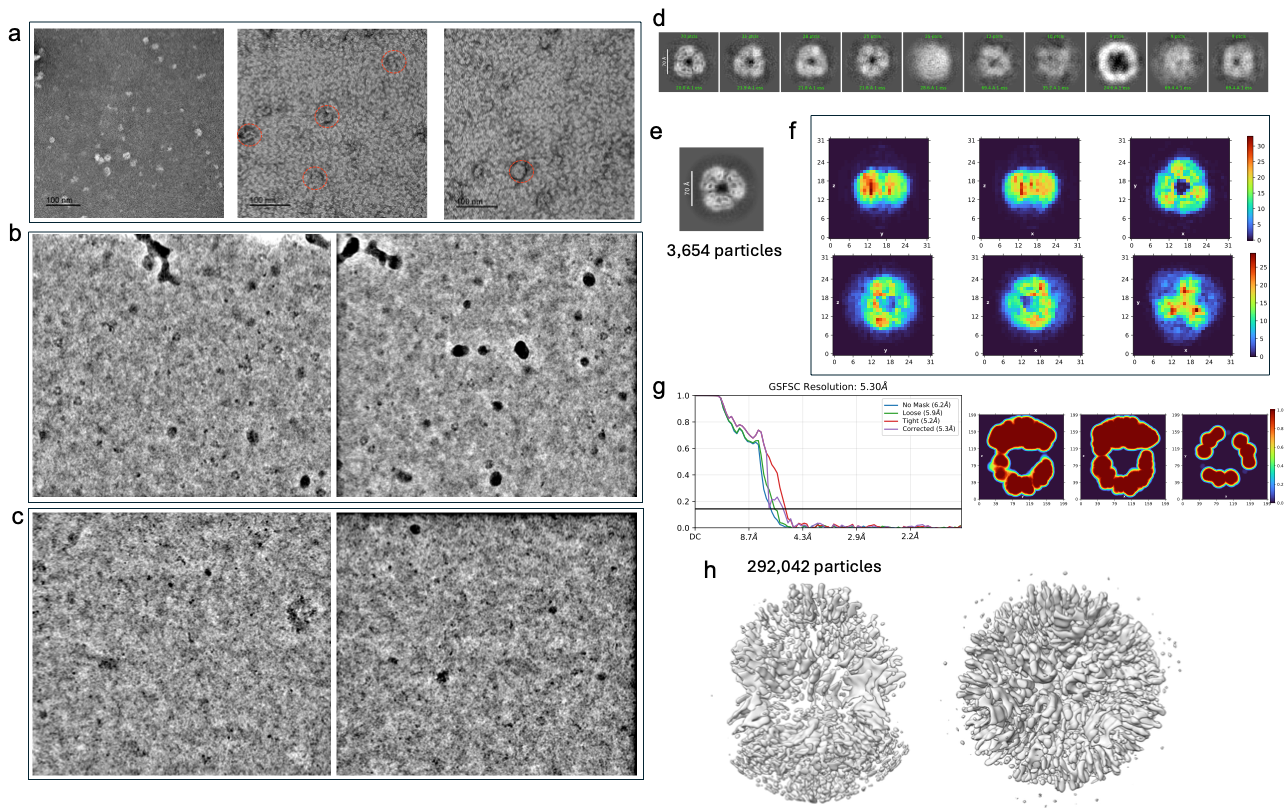

Fig. S18 Cryo-EM analysis of the Lr41-AvrLr41-Lr41NLR^CC^ complex reveals pore-like assemblies. (a) Representative negative-stain electron micrographs of the molar ratio (1:1:1) mixture of Lr41-AvrLr41-Lr41NLR^CC^, showing discrete particles with ring-like features indicated by red dashed circles. TEV protease treated samples were incubated for 1 hr at RT prior to staining. (b) Representative cryo-EM micrographs of Lr41-AvrLr41-Lr41NLR^CC^ mixture, revealing well-defined ring-shaped particles consistent with pore-like assemblies. (c) Representative cryo-EM micrographs of Lr41-AvrLr41 in the absence of Lr41NLR^CC^, showing dispersed particles lacking discernible ring-shaped or pore-like structures as in (b). (d) Examples of manually picked particles from cryo-EM micrographs used for subsequent particle selection. (e) The best 2D-class averages generated following blob-based particle picking guided by the manual picks. (f) Example of *ab initio* 3D reconstructions (C1 symmetry used) generated from selected particles, yielding multiple initial models with central cavities consistent with pore-like architecture. (g) Gold-standard Fourier shell correlation curves and corresponding local resolution maps for the refined reconstruction. (h) Homogeneously refined 3D density map of the Lr41-AvrLr41-Lr41NLR^CC^ complex, shown in two orientations, derived from the full particle dataset and displaying a prominent central pore.

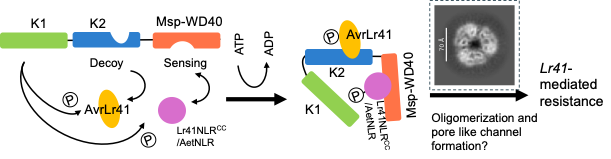

Fig. S19 Simplified working model of Lr41-AvrLr41 recognition and helper-assisted activation. Lr41 comprises a catalytically active kinase (K1), a pseudokinase (K2), and C-terminal MSP-WD40 fusion. AvrLr41 is detected through direct binding to the K2 pseudokinase domain, while the MSP-WD40 region recruits either the truncated helper NLR (Lr41NLR^CC^) or the full-length *Ae. tauschii* helper (AetNLR). Lr41 is proposed to perceive AvrLr41 through a modular kinase architecture in which the pseudokinase domain K2 functions as a decoy, providing a structurally compatible but catalytically inactive surface that binds AvrLr41. Effector engagement at K2 is hypothesized to perturb or stabilize the surrounding domains, repositioning the catalytically active K1 kinase that acts as a guardee within the complex. The C-terminal MSP-WD40 fusion serves as a recruitment platform for the helper NLRs (Lr41NLR^CC^ or AetNLR), enabling formation of a sensor-helper assembly. Effector binding to the K2 decoy domain is proposed to trigger reorganization of K1, K2, and the helper NLR into an activation-competent state required for signal initiation. The assembled complex may subsequently undergo higher-order oligomerisation, forming a resistosome-like structure, as supported by the current observation of pore-shaped particles by cryo-EM, which culminates in Lr41-mediated immune activation.
